# Hyperpolarization-activated cation channel mediated intrinsic plasticity changes underlie the malleability of within cell-type electrophysiological heterogeneity

**DOI:** 10.1101/2025.06.09.658517

**Authors:** Homeira Moradi Chameh, Madeleine Falby, Yvonne Yang, Mandana Movahed, Keon Arbabi, Chaitra Sarathy, Shreejoy J Tripathy, Liang Zhang, Jeremie Lefebvre, Taufik A. Valiante

## Abstract

Within cell-type neuronal electrophysiological, morphological, and transcriptomic heterogeneity is the norm in the brain. Although generally considered a fixed property within cell-types, this heterogeneity is malleable and declines in regions of the human brain that generate seizures. Building off this foundational work we hypothesize that such plasticity of cell-type heterogeneity, specifically its decline, arises from the shared history of neuronal activity that drive intrinsic plasticity mechanisms in a concerted fashion. To explore this hypothesis we study neuronal activity in two model systems: human cortical slice cultures from patients with epilepsy as well as slices from the medial prefrontal cortex (mPFC) and subiculum of rodent kainic acid (KA) model of temporal lobe epilepsy. Biophysical properties and spiking dynamics were characterized using whole-cell patch clamp recordings of layer 2 and layer 3 (L2&3) pyramidal neurons in human slice culture as well as deep layer subicular neurons and layer 5 (L5) mPFC of KA mice. We found a significant decline in biophysical heterogeneity and a reduction in information coding in both the KA and slice culture models. In both these models we found a consistent increase in hyperpolarization-activated cation current (HCN) dependent electrophysiological properties, the blockade of which restored electrophysiological heterogeneity and information coding. Our findings demonstrate that within cell-type heterogeneity is malleable, and despite being a complex distributed network property, can be tuned by a single ionic current. These findings emphasize the plasticity of within cell-type heterogeneity, suggesting the potential for targeted interventions to restore neuronal heterogeneity changes that accompany epilepsy and potentially other neurological and neuropsychiatric diseases.

## 1. Introduction

Heterogeneity of features among entities such as cells, organs, or organisms of the same type is an inherent trait of all biological systems. Over several decades, compelling evidence has shown that this heterogeneity significantly contributes to the stability of physiological functions across all levels of biological structure (Goaillard and Marder 2021; Oliver et al. 2015; Loreau 2000). In the nervous system, heterogeneity is evident not only across different neuronal cell types but also within individual cell types (Cembrowski & Menon, 2018; Cembrowski & Spruston, 2019; Hodge et al., 2019; Scala et al., 2021; Soltesz, 2006; Tasic et al., 2018). This "within-cell-type" heterogeneity arises from variations in morphology and ion channel distributions, determining the unique biophysical properties of neurons that shape how inputs are transformed into outputs. (Hille, 2001; Hodgkin & Huxley, 1952). Heterogeneity in these “intrinsic biophysical properties” of neurons has many computational benefits including improved information coding and learning (Balcioglu et al., 2023; Chelaru & Dragoi, 2008; Gjorgjieva et al., 2016; Luo, 2021; Marsat & Maler, 2010; Padmanabhan & Urban, 2010; Perez-Nieves et al., 2021; Tripathy et al., 2013; Zeldenrust et al., 2021), enhancement of signal detection through resonance (Uzuntarla et al., 2015), functional degeneracy that contributes to network compensation and adaptability (Albantakis et al., 2024; Tononi et al., 1999), and dynamical resilience (Hutt et al., 2023, 2024; Rich et al., 2022).

While the study of neuronal within-cell-type heterogeneity has focused on the computational benefits of intrinsic biophysical heterogeneity, our previous work has provided some of the first evidence for the pathological role of declining intrinsic biophysical heterogeneity in human brain pathology, specifically epilepsy (Rich et al., 2022). We found that L5 pyramidal neurons in epileptogenic regions demonstrated a decline in intrinsic biophysical heterogeneity. Importantly, computational modelling of a spiking network that incorporated this heterogeneity decline was rendered unstable and prone to transitioning into seizure-like states (Rich et al., 2022). More generally, our mathematical analyses have demonstrated that heterogeneity in intrinsic biophysical properties enhances the resilience of neural networks to a wide range of destabilizing factors (Hutt et al., 2023, 2024), suggesting that heterogeneity of biophysical properties is a fundamental aspect of neural circuit function (Cembrowski & Menon, 2018; Cembrowski & Spruston, 2019).

Although within-cell-type heterogeneity is a recognized feature of neuronal networks and likely crucial for normal neuronal function and network stability, the mechanisms by which heterogeneity changes remain poorly understood. *Intrinsic plasticity* - the activity-dependent modulation of ionic conductances, particularly voltage-gated ion channels is a likely candidate mechanism (Beck & Yaari, 2008; Zhang & Linden, 2003). Notably, there is extensive literature that has shown how biophysical properties of neurons are plastic, changing in response to their past activity profiles (Beck & Yaari, 2008; Desai et al., 1999; Turrigiano, 2017; Turrigiano et al., 1994; Zhang & Linden, 2003). Intrinsic plasticity can be *homeostatic*, where a past history of decreased neuronal activity gives rise to increased excitability (or vice versa), or non-homeostatic, where changes in excitability parallel past activity profiles (Turrigiano & Nelson, 2000). In the seminal paper by Turrigiano et al, neurons deprived of activity by blocking voltage-gated sodium channels with tetrodotoxin (TTX), thereby preventing action potential generation, became hyperexcitable (Turrigiano et al., 1994). Importantly, this effect was reversed upon the restoration of activity (Turrigiano et al., 1994). Since these initial observations, subsequent studies extended these findings to cortical regions (Desai et al., 1999), positioning intrinsic plasticity as a fundamental mechanism in memory and learning (Debanne et al., 2019; Zhang & Linden, 2003) on par with synaptic plasticity, and a major contributor to the development of epilepsy (Beck & Yaari, 2008; Timofeev et al., 2014).

Extrapolating these formative findings of intrinsic plasticity from the cellular to the population level suggests that within-cell-type heterogeneity is not a static phenomenon but rather a malleable property. This phenomenon has been demonstrated experimentally, showing that neurons receiving diverse input over time develop distinct transcriptomic profiles (Angelo et al., 2012; Eberwine & Kim, 2015; Park et al., 2014), which in turn would be reflected in the heterogeneity of their biophysical features (Bomkamp et al., 2019; Tripathy et al., 2017). Reciprocally, neurons that share similar patterns of activity would be expected to undergo comparable changes in neuronal excitability (Desai et al., 1999; Mackler et al., 1992; Park et al., 2014; Turrigiano et al., 1994; Tyssowski et al., 2018), leading to a reduction in heterogeneity, a phenomenon we observed in human epileptogenic cortex (Rich et al., 2022). This hypothesis holds regardless of whether the intrinsic plasticity occurs via homeostatic or non-homeostatic mechanisms (Beck & Yaari, 2008; Debanne et al., 2019; Zhang & Linden, 2003). In both cases, neuronal heterogeneity is reduced, either by driving neurons toward a common physiological set point (homeostatic) or by pushing neuronal properties toward extreme but convergent phenotypes (non-homeostatic). For instance, homeostatic plasticity decreases variability by stabilizing firing rates, whereas non-homeostatic plasticity decreases variability by inducing similar long-lasting changes in excitability following sustained activity. Epilepsy is an excellent example of this process, where neurons are driven to have similar activity profiles through hyper-synchrony (Schevon et al., 2012), ultimately resulting in a reduction of intrinsic biophysical heterogeneity (Rich et al., 2022).

To provide a mechanistic understanding of how intrinsic plasticity-mediated biophysical changes contribute to a decline in neuronal heterogeneity, we investigate neurons from human neocortical slice cultures and the rodent kainic acid model of epilepsy. Our rationale for studying human slice cultures is based on the observation that, like neocortical neurons chronically silenced with tetrodotoxin (TTX) (Desai et al., 1999), neurons from slice culture are largely inactive (Johnson & Buonomano, 2007). As such, they represent a low-activity state, in contrast to those participating in the hyper-active epileptiform activity observed in seizure-generating networks (Schevon et al., 2012). Despite the opposite activity levels, both conditions share a common feature: neurons from human slice culture and those generating epileptiform activity have similar past activity profiles. Our human slice culture experiments therefore allow us to dissociate the effects of shared past activity levels from overall activity level on intrinsic biophysical heterogeneity. To complement this work, we also examine neurons from the KA model of epilepsy to assess whether seizure activity, another condition in which neurons share past activity profiles, would similarly lead to a reduction in biophysical heterogeneity.

Using whole-cell patch clamp technique we observed a progressive and profound decline in heterogeneity of intrinsic biophysical properties in cultured human L2&3 neurons over a 72 hour period. In concordance with this observation, L5 pyramidal neurons in the mPFC and deep subicular neurons from the mouse kainic acid model of epilepsy similarly demonstrated a decline in intrinsic biophysical heterogeneity. These demonstrations of reduced heterogeneity were accompanied by decreased variability in spiking dynamics and information content within these populations (Padmanabhan & Urban, 2010; Tripathy et al., 2013). In both conditions, neurons from human slice culture and those generating epileptiform activity, the hyperpolarization-activated cation current (*I*_h_) was observed to increase, which when blocked resulted in an increase in biophysical heterogeneity. These findings, together with our recent computational work (Trotter et al., 2025), provide direct evidence that biophysical heterogeneity is malleable through intrinsic plasticity (Angelo & Margrie, 2011; Eberwine & Kim, 2015; Park et al., 2014; Urban & Tripathy, 2012). They also demonstrate how population heterogeneity can be tuned through modulation of a single ionic current in the cortex (Angelo et al., 2012; Angelo & Margrie, 2011), and highlight the role of shared neuronal activity history in the emergence of pathological states like epilepsy, wherein heterogeneity is aberrantly reduced (Hutt et al., 2023, 2024; Rich et al., 2022; Timofeev et al., 2014).

## 2. Methods

### 2.1 Human ethics and approval

In accordance with the Declaration of Helsinki, institutional approval for this study was received and authorized by the University Health Network Research Ethics Board. The electrophysiological experiments utilized resected brain tissue from eight patients undergoing standard anterior temporal lobectomy (ATL) under general anesthesia with volatile anesthetics (Mansouri et al., 2012). Written informed consent was obtained from each patient prior to surgery, explicitly allowing the use of their brain tissue and human cerebrospinal fluid (hCSF) for research. Additionally, patients consented to the sharing and publication of their anonymized electrophysiological data, along with demographic details including sex, age, duration of seizure history, and antiseizure drug treatments (Table 1).

### 2.2 Animals ethics and approval

All animal experiments were conducted following the guidelines and approval provided by the University Health Network Animal Research Centre. C57BL/6 mice, of both sexes, were utilized in this study. Animals were housed in a controlled vivarium environment maintained at 22°C with regulated humidity and a 12-hour light-dark cycle. Food and water were provided ad libitum.

### 2.3 Kainic acid model of temporal lobe epilepsy

At eight to nine weeks of age, mice were randomly assigned to two experimental groups, receiving stereotaxic injections of either kainic acid or saline into the ventral hippocampus (right dentate gyrus), a region particularly relevant for temporal lobe epilepsy (Zeidler et al., 2018). Mice were anesthetized with 5% isoflurane in 100% oxygen (delivery rate: 1 mL/min) and injected with 100 nL of either kainic acid (20 mM) or sterile saline at coordinates AP 3.6 mm, ML 2.8 mm, DV 2.8 mm from bregma. Infusions were delivered slowly at a rate of 50 nL/min using a motorized syringe (Hamilton Company, USA). Immediately following surgery, mice that received a kainic acid or saline injection were observed for at least 40 minutes where the behavioral severity and durations of their seizures could be recorded according to a modified Racine Scale (Krook-Magnuson & Huntsman, 2005). During this window, all mice that received an injection of kainic acid demonstrated seizures with a range of severities extending between stage 1 and stage 8. In kainic acid or other status models of epilepsy, if animals show extended status seizure-prone, they would be treated with diazepam or other anticonvulsant drug for animal care purposes (Reddy & Kuruba, 2013; Sharma et al., 2018). In general, the severity and/or duration of acute status epilepticus directly influence the latency and incidences of late onset seizures in these models (Rusina et al., 2021). As expected, mice injected with saline did not show any behavioural signs of seizure activity within this postoperative period. To evaluate whether the kainic acid injection was successful in bringing about chronic epilepsy and the manifestation of spontaneous recurrent seizures, kainic acid-injected mice (n = 16) were video-monitored over a 10-day period (Day 26 to Day 35 after surgery) for behavioural signs of seizure activity. Behavioural seizures were initially screened using a Noldus Ethovision XT animal tracking system. Putative seizure activities were automatically detected by a high level of motion activity (5 to 10% above motionless ambient cage behaviors).Video-detected behaviors were then inspected independently by two experimenters to verify behavioral seizures. Severe behavioral seizures, recognized by violent running and jumping with or without falling and four limb clonus, were presented for individual mice (Pitkänen et al., 2017; Velíšková & Velíšek,. 2017).

To characterize the seizure burden for mice receiving an injection of kainic acid, the total number of behavioral seizures, between stage 6 and 8, during the 10-day monitoring period were calculated for a subset of animals (n = 16). Each animal included in this subset demonstrated variable numbers of seizures which can be attributed to expected inter-animal differences in kainic acid sensitivity. The mean number of seizures demonstrated by the subset of animals per day was calculated to be 1.43 over the 10 day monitoring period and comparable to literature values (Zeidler et al., 2018).

### 2.4 In vitro electrophysiology

#### 2.4.1 Slice culture

As previously described (Florez et al., 2013; Moradi Chameh et al., 2021), cortical tissue used for slice cultures was obtained from a surgical resection of a <1 cm³ block from the middle temporal gyrus, a region routinely resected during temporal lobe epilepsy surgery and outside the epileptogenic zone (Moradi Chameh et al., 2023). Immediately after surgical resection, the tissue block was submerged in ice-cold cutting solution containing (in mM): Sucrose 248, KCl 2, MgSO₄·7H₂O 3, CaCl₂·2H₂O 1, NaHCO₃ 26, NaH₂PO₄·H₂O 1.25, and D-glucose 10. This solution, maintained at 4°C and continuously bubbled with hyperoxic carbogen gas (95% O₂ and 5% CO₂), had adjusted pH and osmolarity (7.4 and 300–305 mOsm, respectively).

Transverse brain slices (350 μm thick) were prepared perpendicular to the pial surface using a vibratome (Leica 1200 V) in the same cutting solution, ensuring minimal truncation of pyramidal cell dendrites (Kalmbach et al., 2018; Moradi Chameh et al., 2021). The total duration for transportation and slice preparation was less than 20 minutes (Moradi Chameh et al., 2021). Slices were then incubated at 34°C for 30 minutes in recovery chambers containing artificial cerebrospinal fluid (aCSF; in mM: NaCl 123, KCl 4, CaCl₂·2H₂O 1.5, MgSO₄·7H₂O 1.3, NaHCO₃ 26, NaH₂PO₄·H₂O 1.2, and D-glucose 10), continuously carbogenated (95% O₂ and 5% CO₂), with maintained pH (∼7.4) and osmolarity (300–305 mOsm). Following this incubation, slices were stored at room temperature (22–23°C) in carbogenated aCSF for at least an additional 30 minutes.

Subsequently, slices were transferred onto uncoated Millicell-CM culture inserts (30 mm diameter, 0.4 µm pore size, Millipore) and cultured in media containing 48% DMEM/F-12 (Gibco), 48% Neurobasal (Gibco), 1x N-2 (Gibco), 1x B-27 (Gibco), 1x Glutamax (Gibco), 1x NEAA (Gibco), and 20 mM HEPES (Gibco). After one hour, slices were moved into plates containing sterile-filtered human cerebrospinal fluid (hCSF), previously centrifuged at 4000 rpm at 4°C for 10 minutes and stored at –80°C (Bak et al., 2024; Schwarz et al., 2019). Plates were incubated at 37°C in 5% CO₂ and 100% humidity. All tools and equipment used for slice culture and preparation were sterilized in advance using UV light, heat, or ethanol, and slice preparation was performed under sterile conditions.

#### 2.4.2 Subiculum and mPFC slice preparation

Between 35 and 42 days following KA or saline injection, mice underwent in vitro whole-cell patch-clamp experiments targeting the deep pyramidal layer of the subiculum or L5 mPFC. Mice were deeply anesthetized with isoflurane vapor (>5%), decapitated, and brains quickly removed and submerged in ice-cold cutting solution saturated with carbogen gas (95% O₂, 5% CO₂). The cutting solution contained (in mM): Sucrose 160, D-glucose 7.25, NaHCO₃ 28, HEPES 20, KCl 2.5, NaH₂PO₄·H₂O 1.25, Sodium Ascorbate 3, Sodium Pyruvate 3, MgCl₂·6H₂O 7.5, CaCl₂·2H₂O 1 (pH adjusted to 7.4, osmolarity 295–305 mOsm (Beaulieu-Laroche et al., 2018).

Slices (350 μm) ipsilateral to the injection site were prepared using a Leica Vibratome (VT1200S). Ventral hippocampal slices (subiculum) were cut transversely at depths of 1500–2500 μm from the ventral surface, while coronal slices of mPFC were cut from the rostral brain end. Slices were transferred to a recovery chamber containing aerated aCSF composed of (in mM): NaCl 120, D-glucose 11, NaHCO₃ 25, KCl 3, NaH₂PO₄·H₂O 1.25, sodium ascorbate 1, sodium pyruvate 3, MgCl₂·6H₂O 1.5, and CaCl₂·2H₂O 1.5 (pH ∼7.4, osmolarity 295–305 mOsm) (Beaulieu-Laroche et al., 2018). Slices were incubated at either 32°C (subiculum) or 34°C (mPFC) for 30 minutes and subsequently transferred to room temperature (22–23°C) for at least 1 hour before recording. Slices remained viable for approximately 5 hours.

### 2.5 Whole-cell patch clamp recording and data acquisition

For electrophysiological recordings, slices were transferred to a recording chamber mounted on a fixed-stage upright microscope. L5 pyramidal neurons of the mPFC were visualized using an Olympus BX51WI upright microscope equipped with differential interference contrast (DIC) optics and an infrared Retiga ELECTRO CCD camera (IR-1000, Teledyne Photometrics). Deep pyramidal neurons of the subiculum were visualized using a Nikon Eclipse FN1 microscope with DIC optics, an IR-CCD camera (IR-1000, MTI), and a ×40 water immersion objective. Slices were continuously perfused with standard aCSF at 4 ml/min at room temperature. L5 of the prelimbic (PrL) cortex in the mPFC was easily distinguishable, located at a perpendicular distance of 300–500 µm from the midline. The PrL cortex, positioned along the midline, is bordered dorsally by the anterior cingulate cortex and ventrally by the infralimbic (IL) cortex. In the subiculum, recordings were conducted in the deep cell layers proximal to CA1. Pipettes (3–6 MΩ resistance), filled with intracellular solution containing (in mM): 135 K-gluconate, 10 NaCl, 10 HEPES, 1 MgCl₂, 2 Na₂ATP, and 0.3 GTP (pH 7.3, 290–309 mOsm), were pulled from thin-wall borosilicate glass (World Precision Instruments, USA) using a vertical puller (PC-10, Narishige, Japan).Whole-cell patch-clamp recordings were obtained using a Multiclamp 700B amplifier, and data acquisition was performed with 1440A and 1550B digitizers using pClamp software (version 11; Axon Instruments, Molecular Devices, USA) at a sampling rate of 20 kHz. Passive and active electrophysiological properties were assessed under current-clamp conditions, with synaptic activity blocked by including GABAergic and glutamatergic receptor antagonists in the perfusion aCSF. Specifically, bicuculline (10 μM) and CGP (10 μM) were used to block GABA-A and GABA-B receptors, respectively, while CNQX (10 μM) and APV (20 μM) were employed to block AMPA and NMDA glutamate receptors. Cells were subjected to hyperpolarizing and depolarizing current pulses ranging from -250 pA to +300 pA in ±50 pA increments, each lasting 600 ms. When indicated, ZD-7288 (10 μM), an hyperpolarization-activated cyclic nucleotide-gated (HCN) channel blocker (HCN), was applied. Access resistance (Ra) was continuously monitored and maintained within an acceptable range of 16–32 MΩ.

Heterogeneity in input-output transformation can also be captured by the spiking dynamics of neurons within a population (Padmanabhan & Urban, 2010). Intuitively, a population of neurons that each generate identical firing patterns to a frozen white noise stimulus convey less information (lower entropy) about the stimulus than a population of neurons where each neuron generates a distinct spike train (higher entropy) (Padmanabhan & Urban, 2010). In a homogeneous population, the spike train from a single neuron captures the ensemble activity, whereas if all the neurons are biophysically diverse, then the output of every neuron is required to capture the ensemble response. To quantify the heterogeneity in spiking dynamics, neurons were stimulated with a 10-trial (inter-trial interval = 20 s) current injection stimulus of frozen white noise of 2.5 ms duration convolved with a 3-ms square function. A consistent level of DC current (Inibhunu et al., 2023; Padmanabhan & Urban, 2010) was added to the noisy current input (40 pA), with incremental changes in DC amplitude to induce spiking at 6-8Hz (ie. theta frequency), a dominant rhythm in both the human and rodent brain (Groppe et al., 2013). Peaks in the voltage trace exceeding 0 mV were identified as action potentials (Moradi Chameh et al., 2021).

### 2.6 Intrinsic biophysical feature analysis

Whole-cell patch-clamp recording data were analyzed using pClamp/Clampfit (version 11), GraphPad Prism (version 9.5.0), and a custom Python-based script. Passive membrane properties calculated for each neuron included resting membrane potential (RMP), membrane potential (MP), membrane time constant (MTC), and input resistance (IR). RMP was defined as the neuron’s membrane voltage at rest immediately after membrane rupture, without current injection (I = 0 pA). MP was calculated as the average membrane voltage measured 5 minutes after cell rupture and before current injection commenced. The MTC, representing the time required for the neuron’s voltage potential to reach approximately 63% of its RMP, was determined by fitting a standard exponential function to the voltage response measured between 10 ms and 60 ms after a -50 pA hyperpolarizing current injection lasting 600 ms. IR reflects the number of open membrane channels and was calculated by measuring the voltage deflection in response to hyperpolarizing current injections (-200 to -50 pA, step size 50 pA, 600 ms duration). A linear regression analysis of voltage responses versus current injection amplitude provided the slope, corresponding to the input resistance. Active electrophysiological properties calculated included action potential threshold, rheobase, delay to first spike, time to peak, distance to threshold (DTT), sag voltage amplitude, average firing rate, and interspike interval (ISI). These parameters were assessed using hyperpolarizing and depolarizing square-pulse current injections ranging from -250 pA to +350 pA in increments of 50 pA, each with a duration of 600 ms.

Threshold was defined as the minimum membrane potential (mV) required to initiate an action potential. Rheobase was the minimum depolarizing current amplitude (pA) necessary to trigger an action potential. Delay to first spike was measured as the latency (ms) from the onset of depolarizing current injection to the initiation of an action potential, time to peak was defined as the time (ms) from the baseline membrane potential to the maximum voltage of the action potential. DTT represents the voltage difference (mV) from the resting potential to the threshold for spike initiation. Sag voltage amplitude was quantified as the absolute difference between the voltage trough elicited by a -250 pA hyperpolarizing current injection and the steady-state voltage before the end of the current pulse. The average FI curve was generated by calculating the instantaneous firing frequency at each spike for depolarizing current injections (50–250 pA, in 50 pA increments, 600 ms duration) and then averaging these frequencies across the cell population. The ISI, inversely related to firing rate, represented the duration between consecutive action potentials elicited by depolarizing currents. Additionally, the local variation in inter-spike interval (LV-ISI) was calculated instead of the traditional coefficient of variation (CV ISI), as CV ISI values can be biased by fluctuations in firing rate, leading to increased scatter in ISI measurements (Shinomoto et al., 2009). The local variation (Lv) is defined as follows (Equation 1): if *T_i_* represents the i-th inter-spike interval and 𝑛 represents the total number of intervals within a trace, then Lv is calculated by

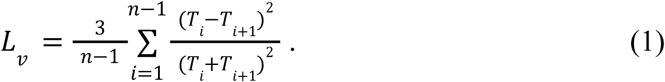

### 2.7 Analysis of neuronal spiking variability and population coding under white noise stimulation

White noise analysis was performed on single-cell voltage recordings obtained through injected white noise current, following the methodology described by Padmanabhan et al (Padmanabhan & Urban, 2010). Principal Component Analysis (PCA) was conducted separately for each treatment group across all sweeps (n = 10 per cell) from each cell (n = 36 for kainic acid model; n = 33 for saline; for slice culture: n = 13 at 0 hr, n = 16 at 24 hr, n = 13 at 48 hr, n = 10 at 72 hr; n = 9 before ZD-7288 application, n = 9 after ZD-7288 application). The PCA involved calculating the covariance matrix for all sweeps from each cell within the respective treatment groups and ranking the eigenvectors according to their eigenvalues to identify principal components. These principal components served as features for classifying the originating cell using the k-Nearest-Neighbors (k-NN) algorithm (Padmanabhan & Urban, 2010). Prediction accuracy was measured as the proportion of sweeps in the test dataset correctly assigned to their originating cell. Both the parameter k and the number of principal components included were systematically varied to assess their impact on prediction accuracy. The k-NN classifier was trained on (100 - T)% of the data and tested on the remaining unseen T%, where T varied among (T∈{10, 20, 30, 35, 40}). This training-testing process was repeated five times with randomly selected training sets to minimize bias.

Entropy was calculated to measure the informational content that could be encoded by neuronal populations. To compute the entropy for a given population of n cells, voltage recordings were partitioned into 5 ms bins and then binarized, assigning a value of 1 if a bin contained at least one spike, and 0 otherwise (Padmanabhan & Urban, 2010). Through this binarization, each combination

𝐶_𝑖_ of *n* cells generated an n-bit vector for each time bin. The event space WCi was defined as the set of all unique *n*-bit encodings observed in the sweeps of n-cell recordings, with the probability 𝑃(𝑤 ∈ 𝑊_𝐶_i) representing the likelihood of encoding 𝑊𝐶𝑖 occurring. Shannon entropy, measured in bits due to the logarithm base 2, was calculated as:

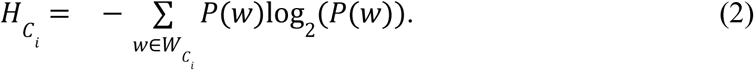

The average entropy was computed across various combinations Ci of selecting n cells from a total of N cells. In cases with large numbers of cells (e.g., N = 36 for kainic acid model and N = 33 for saline), entropy estimation was performed for combinations of up to 1000 cells, as exact computation became computationally impractical (33 choose 17=1166803110).

Additionally, the cardinality of 𝑊_𝐶𝑖_ represents the number of unique encodings recorded. Due to the sparse nature of spiking activity in response to white noise injections, the encoding consisting entirely of zeros often occurred disproportionately, reducing distribution uniformity and thus lowering entropy. Therefore, bin size was optimized to minimize the frequency of all-zero encodings (maximizing entropy), while ensuring no bin contained more than one spike.

### 2.8 Single-nucleus RNA sequencing (snRNA-seq) analysis of intrahippocampal kainic acid-induced mouse model of temporal lobe epilepsy

At 12 weeks following the injection of either kainic acid or saline targeted to the right ventral hippocampus (dentate gyrus), mice were prepared for single-nucleus RNA sequencing (snRNA-seq). Mice were deeply anesthetized via inhalation of isoflurane vapor (>5%) for approximately 20 seconds, with anesthesia confirmed by the absence of a response to a toe pinch (loss of pedal withdrawal reflex). Subsequently, mice underwent cardiac perfusion with ice-cold 1X phosphate-buffered saline (PBS). Brain tissues were promptly dissected, snap-frozen in liquid nitrogen, and stored at -80°C until further processing. Three biological replicates per experimental group were prepared to enable robust differential gene expression analysis. snRNA-seq was performed using a 10X Genomics Chromium X Controller, targeting approximately 10,000 nuclei per sample with a sequencing depth of 60,000 reads per nucleus.

Raw gene expression matrices were generated using the 10X Genomics platform and imported into Seurat (Satija et al., 2015) through the Read10X function to create Seurat objects. Quality control was applied by filtering out nuclei with fewer than 2000 detected features or greater than 10% mitochondrial gene content, using the subset function in Seurat. This filtering step ensured the removal of low-quality cells and potential cellular debris. Subsequently, highly variable genes were identified, and principal component analysis (PCA) was performed. Clustering analysis utilized the FindNeighbors and FindClusters functions in Seurat, with a clustering resolution set to 0.5. For dimensionality reduction and visualization, uniform manifold approximation and projection (UMAP) was conducted with the following parameters: dimensions = 1:30, n.epochs = 500, min.dist = 0.3, and spread = 1. Variability in gene expression within each cell type and experimental group was quantified by calculating the standard deviation (SD) of normalized gene expression, with lower SD indicating reduced transcriptomic heterogeneity. Statistical significance of differential expression was assessed using non-parametric tests.

### 2.9 Statistical analyses

All statistical analyses of intrinsic biophysical properties were performed using GraphPad Prism (version 9.5.0). Normality was assessed using the D’Agostino & Pearson, Anderson-Darling, Shapiro-Wilk, and Kolmogorov-Smirnov tests. Properties that failed these tests were considered non-normally distributed. Statistical comparisons were conducted between the following experimental groups: (1) deep subiculum pyramidal neurons under seizure-prone (kainic acid injection) versus non-seizure-prone (control) conditions (saline injection); (2) L5 mPFC pyramidal neurons under seizure-prone versus non-seizure-prone conditions; (3) L2&3 pyramidal neurons from human tissue cultured slices at multiple time points across four days (comparisons: 0 vs. 24 hours, 0 vs. 48 hours, 0 vs. 72 hours, 24 vs. 48 hours, 48 vs. 72 hours, and 24 vs. 72 hours); and (4) L5 somatosensory pyramidal neurons before and after application of ZD-7288, an HCN channel blocker. To initially determine if intrinsic biophysical properties exhibited significant differences in means across various comparison groups, either a parametric Welch’s unpaired t-test or a non-parametric Mann-Whitney unpaired test was applied, depending on the normality of each property. Subsequently, differences in variability for each property between groups were evaluated. For normally distributed properties, variability differences were assessed using the two-sample coefficient of variation test (CV2 TEST function) based on the Krishnamoorthy and Lee method, implemented via the Real Statistics Resource Pack Excel package (https://real-statistics.com/) (Krishnamoorthy & Lee, 2014). Non-normally distributed data were first transformed using log10 or square root (SQRT) transformations, followed by normality tests to confirm successful normalization before performing the CV2 test (Buzsáki & Mizuseki, 2014).

## 3. Results

### 3.1 Human slice culture

Our preliminary investigations (Rich et al., 2022) revealed a reduction of intrinsic biophysical heterogeneity in acutely resected human neocortical tissue from regions identified electrophysiologically as hyperactive, based on the epileptiform activity observed in intraoperative electrocorticographic recordings (Greiner et al., 2016). The observed reduction in heterogeneity among neurons that share a history of activity, particularly in seizure states, motivated our current investigation into whether a similar phenomenon occurs in other conditions marked by correlated patterns of activity. Specifically, we used slice culture experiments, in which neurons experience a shared history of activity deprivation, to determine whether mechanisms of intrinsic plasticity similarly drive a loss of intrinsic biophysical heterogeneity.

The seminal work by Desai et al. (1999) provided compelling experimental evidence for such activity-deprived regulation of intrinsic neuronal excitability (Desai et al., 1999). They showed that by applying TTX to block action potential generation, cortical pyramidal neurons undergo a shift in their intrinsic properties, most notably, an increase in excitability via selective modulation of sodium and potassium currents (Desai et al., 1999). Therefore, neurons in slice culture by sharing activity patterns through quiescence (Johnson & Buonomano, 2007), should demonstrate a decline in biophysical heterogeneity. To explore this question, we targeted L2&3 pyramidal neurons in cultured human cortical slices to complement our initial findings in L5 pyramidal neurons (Rich et al., 2022), aiming to test whether the phenomenon extends across cortical layers, as suggested by our transcriptomic analyses (Moradi Chameh et al., 2023).

We performed whole-cell patch-clamp recordings from L2&3 pyramidal neurons in acute and cultured neocortical brain slices derived from the MTG of eight individuals who underwent resective surgery for pharmacoresistant epilepsy (see **Supplementary Table 1** for patient details). Previous work has shown that these neurons in acute slices exhibit substantial electrophysiological heterogeneity (Berg et al., 2020; Moradi Chameh et al., 2021). The MTG is a well-characterized region commonly used as a source of non-epileptogenic tissue in both electrophysiological and transcriptomic studies of human cortical neurons (Beaulieu-Laroche et al., 2018; Hodge et al., 2019; Kalmbach et al., 2018; Moradi Chameh et al., 2021). To track the changes in biophysical heterogeneity over time, we measured both passive and active electrophysiological properties in L2&3 pyramidal neurons acutely (0 Hr) and at 24, 48, and 72 hours post-culturing (n = 41, n = 24, n = 14, and n = 10 neurons, respectively). Representative voltage traces from hyperpolarizing current steps, which were used to calculate passive membrane properties, are shown in **Fig. 1A**.

**Figure 1.**
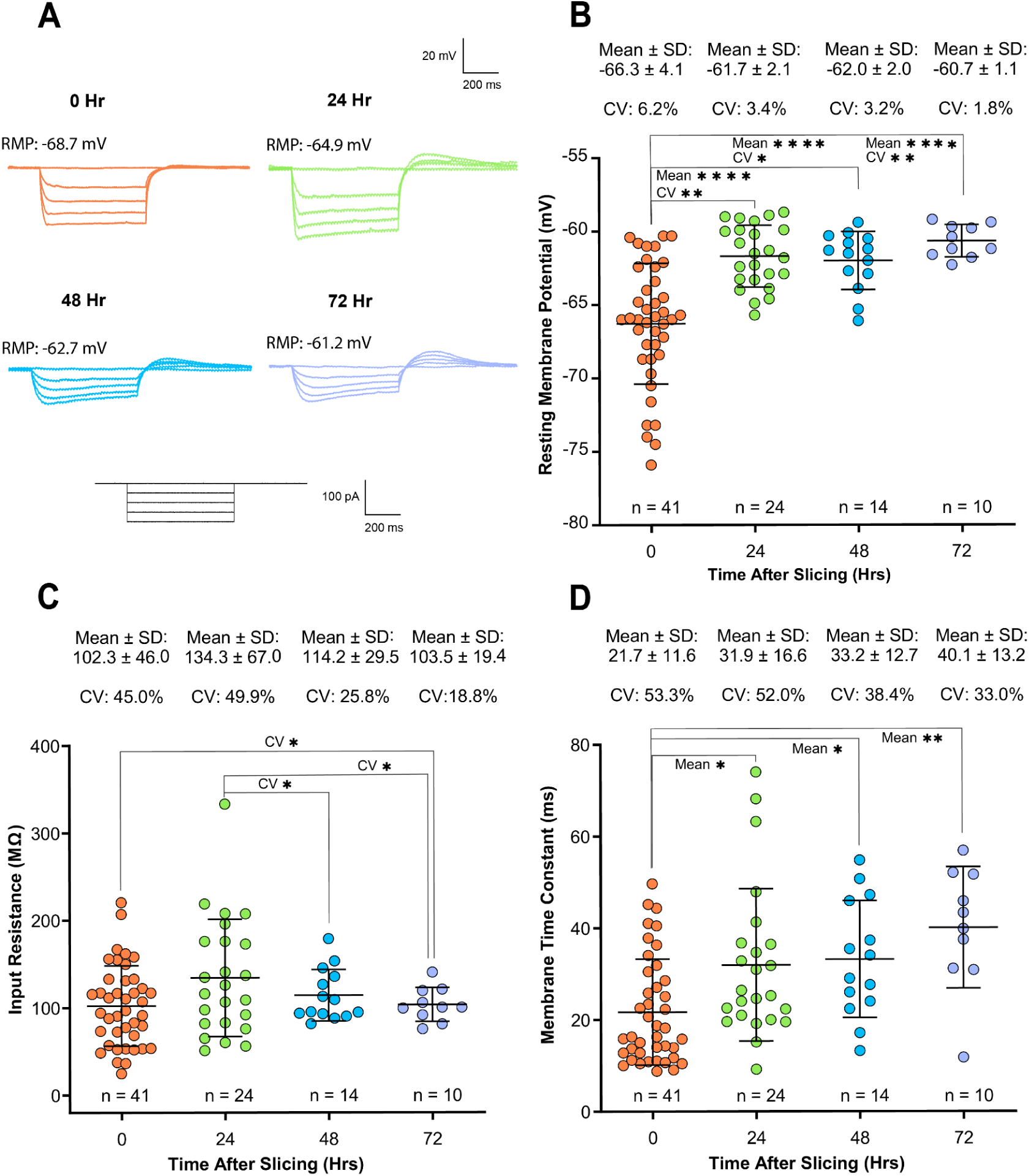
| Heterogeneity decline and increased excitability of human cortical neurons in slice culture: **(A)** Representative voltage traces in response to hyperpolarizing current injections used to calculate passive membrane properties in each condition (see Methods). **(B)** Cultured neurons exhibited a significant depolarization of RMP compared to acute recordings, accompanied by a progressive reduction in RMP variability. The coefficient of variation (CV) for RMP significantly declined over time (**CV:** acute vs. 24 Hr: p = 0.0011; acute vs. 48 Hr: p = 0.0149; acute vs. 72 Hr: p = 0.0018). Coefficient of variation comparisons were performed using a two-sample CV test. Mean comparisons similarly revealed highly significant differences across all post-culture time points (**Mean:** acute vs. 24 Hr: p < 0.0001; acute vs. 48 Hr: p < 0.0001; acute vs. 72 Hr: p < 0.0001; 48 Hr vs. 72 Hr: p = 0.0465; one-way ANOVA with Tukey’s multiple comparisons test). **(C)** Input resistance variability significantly decreased at 72 hours compared to acute conditions (**CV:** acute vs. 72 Hr: p = 0.0323), and was also significantly reduced at both 48 and 72 hours compared to 24-hour cultures (**CV:** 24 Hr vs. 48 Hr: p = 0.0392; 24 Hr vs. 72 Hr: p = 0.0185). No significant differences were observed in mean input resistance across time points (**Mean:** acute vs. 24 Hr: p = 0.0594, acute vs. 48 Hr: p = 0.8579; acute vs. 72 Hr: p = 0.9999). **(D)** No significant changes in the variability of the membrane time constant were observed (**CV:** acute vs. 24 Hr: p = 0.3668; acute vs. 48 Hr: p = 0.1648; acute vs. 72 Hr: p = 0.2816). However, the mean membrane time constant significantly increased following culturing (**Mean:** acute vs. 24 Hr: p = 0.0459; acute vs. 48 Hr: p = 0.0239; acute vs. 72 Hr: p = 0.0014; Kruskal–Wallis test with Dunn’s post hoc comparison).

#### 3.1.1 Heterogeneity decline in passive biophysical properties accompanies activity deprivation in human slice culture

Changes in intrinsic biophysical heterogeneity were quantified using the coefficient of variation (CV) for each measured parameter, and CV values among groups were compared using two-sample coefficients of variation tests (CV2 tests) (Rich et al., 2022). A progressive decline in RMP variability was observed from acute to 72 hours post-culturing (acute CV: 6.2%; 24 Hr CV: 3.4%; 48 Hr CV: 3.2%; 72 Hr CV: 1.8%; **Fig. 1B**), with significant reductions correlated to longer culture durations (acute vs. 24 Hr: p = 0.0011; acute vs. 48 Hr: p = 0.0149; acute vs. 72 Hr: p = 0.0018). Similarly, input resistance variability significantly decreased at 72 hours compared to acute conditions (acute CV: 45.0%; 72 Hr CV: 18.8%; p = 0.0323), and was significantly lower at both 48 and 72 hours compared to 24-hour cultures (24 Hr CV: 49.9%; 48 Hr CV: 25.8%, p = 0.0392; 72 Hr CV: 18.8%, p = 0.0185; **Fig. 1C**). Lastly, although membrane time constant variability showed a declining trend post-culture, it did not reach statistical significance (acute CV: 53.3%; 24 Hr CV: 52.0%; 48 Hr CV: 38.4%; 72 Hr CV: 33.0%; acute vs. 24 Hr: p = 0.3668; acute vs. 48 Hr: p = 0.1648; acute vs. 72 Hr: p = 0.2816; **Fig. 1D**).

These changes in heterogeneity were also accompanied by homeostatic changes in excitability as previously observed in cortical neurons deprived of activity (Desai et al., 1999). We found that after culturing the slices for three consecutive days, the neurons had substantially more depolarized RMPs compared to when they were recorded acutely (acute: -66.3 ± 4.1 mV; 24 Hr: -61.7 ± 2.1 mV; 48 Hr: -62.0 ± 2.0 mV; 72 Hr: -60.7 ± 1.1 mV; **Fig. 1B**). Additionally, the RMP displayed a significant positive correlation with the time of culturing (acute compared to 24 HFr: p < 0.0001; acute compared to 48 Hr: p = 0.0002; acute compared to 72 Hr: p < 0.0001; One-way ANOVA with Tukey’s multiple comparisons test), likely due to enhancement of *I*_h_ currents, consistent with previous studies (Ting et al., 2018). Additionally, input resistance remained stable following culturing compared to acute conditions (acute: 102.3 ± 46.0 MΩ; 24 Hr: 134.3 ± 67.0 MΩ; 48 Hr: 114.2 ± 29.5 MΩ; 72 Hr: 103.5 ± 19.4 MΩ; **Fig. 1C**), aligning with previous findings (Schwarz et al., 2019; Ting et al., 2018). Moreover, there was little to no relationship between input resistance and time post-slicing (acute vs. 24 Hr: p = 0.0594; acute vs. 48 Hr: p = 0.8589; acute vs. 72 Hr: p = 0.9999; One-way ANOVA with Tukey’s multiple comparisons test). The membrane time constant significantly increased following culturing compared to acute (acute: 21.7 ± 11.6 ms; 24 Hr: 31.9 ± 16.6 ms, p = 0.0459; 48 Hr: 33.2 ± 12.7 ms, p = 0.0239; 72 Hr: 40.1 ± 13.2 ms, p = 0.0014; Kruskal–Wallis test with Dunn’s post hoc comparison; **Fig. 1D**).

#### 3.1.2 Heterogeneity decline in active biophysical properties accompanies activity deprivation in human slice culture

We extended our analysis of passive biophysical properties to explore whether similar decreases in variability are observed in spiking behavior. We observed a consistent decline in the variability of multiple electrophysiological properties with increased time in culture. Example voltage traces in response to depolarizing and hyperpolarizing current steps, used to assess active membrane properties, are shown in **Fig. 2A**. Rheobase variability (CV) decreased from 50.5% (acute) to 27% (72 Hr), with significant differences between acute and 72 Hr (p = 0.0330), and between 24 Hr (63.7%) and 72 Hr (p = 0.0145; **Fig. 2B**). Action potential amplitude variability also declined (acute: 20.1%; 72 Hr: 7.9%), with significant reductions at 48 Hr (p = 0.0479), 72 Hr (p = 0.0109), and between 24 and 72 Hr (p = 0.0251; **Fig. 2C**). Sag voltage amplitude variability significantly decreased from acute (65.1%) to 48 Hr (49.6%, p = 0.0240; **Fig. 2D**). Firing rate variability declined significantly at specific current injections: 100 pA (acute vs. 24 hr: p = 0.0138), 200 pA (acute vs. 72 Hr: p = 0.0052; 24 Hr vs. 72 Hr: p = 0.0031; 48 Hr vs. 72 hr: p = 0.0039), and 250 pA (acute vs. 72 Hr: p = 0.0065; 48 Hr vs. 72 Hr: p = 0.0039), as illustrated by bar graphs highlighting the observed loss of variability at these current levels **(Fig. 2E)**.

**Figure 2.**
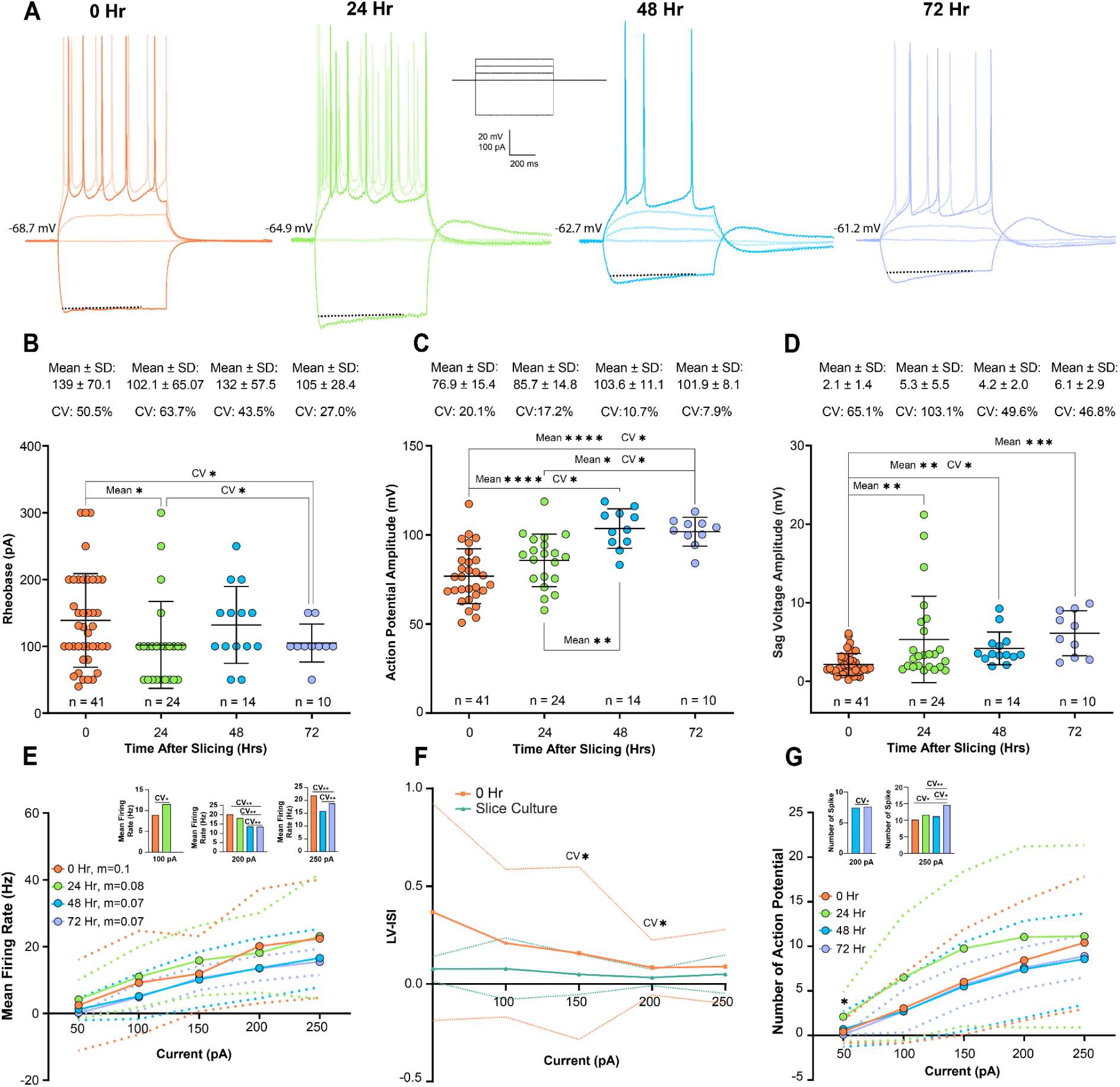
| Changes in heterogeneity and mean of active electrophysiological properties in human cortical layer 2&3 pyramidal cells during slice culture: **(A)** Representative voltage traces in response to depolarizing and hyperpolarizing current injections used to assess active membrane properties across conditions (see Methods). **(B–G)** A consistent reduction in the heterogeneity (coefficient of variation, CV) of active properties was observed with increasing time in culture. **(B)** Rheobase variability significantly decreased between acute and 72 Hr (p = 0.0330) and between 24 Hr and 72 Hr (p = 0.0145). Mean rheobase was also significantly reduced at 24 Hr compared to acute (p = 0.0492; Kruskal–Wallis with Dunn’s post hoc test). **(C)** Action potential amplitude variability (at 150 pA) decreased post-culturing (acute vs. 48 Hr: p = 0.0479; acute vs. 72 Hr: p = 0.0193; 24 Hr vs. 72 Hr: p = 0.0251). In parallel, mean amplitude significantly increased at 48 Hr and 72 Hr compared to acute (both p < 0.0001), and 24 Hr was significantly lower than 48 Hr (p = 0.0054) and 72 Hr (p = 0.0185; one-way ANOVA with Tukey’s test). **(D)** Sag voltage amplitude variability significantly declined at 48 Hr compared to acute (p = 0.0240). Mean sag amplitude progressively increased over time (acute vs. 24 Hr: p = 0.0069; acute vs. 48 Hr: p = 0.0020; acute vs. 72 Hr: p = 0.0002; Kruskal–Wallis with Dunn’s post hoc). **(E)** Firing rate variability decreased consistently at specific current injections (100 pA, acute vs. 24 Hr: p = 0.0138; 200 pA, acute vs. 72 Hr: p = 0.0052; 24 Hr vs. 72 Hr: p = 0.0031; 48 Hr vs. 72 Hr: p = 0.0039; 250 pA, acute vs. 72 Hr: p = 0.0065; 48 Hr vs. 72 Hr: p = 0.0039), as illustrated by the bar graphs at these currents, while the mean firing rate remained unchanged across time points. **(F)** LV-ISI, a measure of spike timing variability, significantly decreased at 100 pA and 200 pA in cultured slices (p = 0.0313 and p = 0.0312, respectively), indicating more regular firing over time. Mean LV-ISI did not differ across conditions. **(G)** The number of action potentials showed stable mean values across time points, except for a significant reduction at 50 pA between 24 Hr and 72 Hr (p = 0.0192; two-way ANOVA with Tukey’s multiple comparisons test). However, heterogeneity in the number of action potentials declined significantly over time. At 200 pA, CV decreased from 48 Hr to 72 Hr (p = 0.0496), and at 250 pA, 72 Hr cultures showed significantly reduced variability compared to acute, 24 Hr, and 48 Hr (p = 0.0178, p = 0.0082, and p = 0.0314, respectively. As illustrated by the bar graphs at these currents). The two-sample coefficient of variation test was used to compare CV between experimental groups.

Spike timing variability, measured by LV-ISI, significantly decreased at both 100 pA (p = 0.0313) and 200 pA (p = 0.0312; **Fig. 2F**), indicating increased temporal regularity. Similarly, variability in the number of action potentials significantly decreased at 200 pA between 48 and 72 Hr (p = 0.0496), and at 250 pA between 72 Hr and all earlier time points (acute: p = 0.0178; 24 Hr: p = 0.0082; 48 Hr: p = 0.0314; as illustrated by the bar graph **Fig. 2G**). Variability comparisons were made using a two-sample coefficient of variation test.

In terms of mean values, rheobase significantly decreased at 24 Hr vs. acute (139 ± 70.1 pA vs. 102.1 ± 65.0 pA, p = 0.0492), with no significant differences at 48 or 72 Hr (Kruskal–Wallis with Dunn’s post hoc; **Fig. 2B)**. Action potential amplitude increased significantly at 48 Hr (103.6 ± 11.8 mV) and 72 Hr (101.9 ± 8.1 mV) vs. acute (76.9 ± 15.4 mV; both p < 0.0001), and was also higher than at 24 Hr (85.7 ± 14.8 mV; p = 0.0054 and p = 0.0185 respectively; one-way ANOVA with Tukey’s test; **Fig. 2C**). Sag voltage amplitude increased progressively over time (acute: 2.1 ± 1.4 mV; 24 Hr: 5.3 ± 5.5 mV; 48 Hr: 4.2 ± 2.1 mV; 72 Hr: 6.12 ± 2.9 mV), with significant differences from acute at all time points (p = 0.0069, 0.0020, and 0.0002 respectively; **Fig. 2D**), consistent with increased HCN channel activity. Mean firing rate and LV-ISI were unchanged across time (**Figs. 2E & 2F**), and no significant differences were found in other active properties, such as DTT, delay to first spike, or threshold (**Supplementary** Fig. 1A–C). The mean number of action potentials remained stable across time, except at 50 pA where a reduction was observed between 24 Hr and 72 Hr (2.1 ± 2.8 vs. 0.1 ± 0.32; p = 0.0192; **Fig. 2G**).

Our analysis of changes in passive and active membrane properties reveal a progressive loss of intrinsic biophysical heterogeneity in important excitability parameters of cortical pyramidal neurons over 72 hours in culture, alongside homeostatic changes in excitability (Desai et al., 1999). These reductions in population-level biophysical heterogeneity suggest that in-vivo spontaneous activity supports a richer distribution of excitability states, while culturing, effectively a low-activity state, leads to decrease in the biophysical heterogeneity. In parallel, neurons became progressively more depolarized and exhibited longer membrane time constants, consistent with homeostatic compensation mechanisms previously observed under activity blockade (Desai et al., 1999). Together, these suggest that intrinsic variability is dynamically modulated by ongoing population activity, and lost when that activity is reduced or silenced.

### 3.2 Epilepsy-related decline in biophysical heterogeneity of passive and active properties in the kainic acid mouse model

An alternative approach for the imposition of shared history of activity, is to subject neurons to the periodic correlated inputs like those that occur in epilepsy (Schevon et al., 2012). Such correlated output from the epileptogenic zone, generate similar activity profiles in downstream regions, which should result in heterogeneity decline (Angelo et al., 2012; Eberwine & Kim, 2015; J et al., 2015; Mackler et al., 1992; Park et al., 2014). We chose the hippocampal KA model of epilepsy as a naturalistic and well established model approach to study changes in neuronal heterogeneity arising from correlated inputs (Pitkänen et al., 2017).

L5 pyramidal neurons in the mPFC and deep-layer subicular neurons are monosypatically connected to the hippocampus making them direct recipients of the the correlated activity arising from the epileptogenic hippocampus (Choy et al., 2022; Drexel et al., 2012; Fei et al., 2022; Gill et al., 2016; Stafstrom, 2012). These regions have been shown to progressively become engulfed into the epileptogenic network, resulting in the brain-wide propagation (generalization) of seizure activity (Choy et al., 2022; Fei et al., 2022). We hypothesize that this transformation from focal to generalized seizures is accompanied by a decline in heterogeneity in the mPFC and subiculum reducing the network’s capacity to desynchronize activity (Hutt et al., 2023, 2024; Padmanabhan & Urban, 2010; Rich et al., 2022), thereby enabling the spread of correlated, seizure-like discharges throughout the brain.

#### 3.2.1 Decline in heterogeneity of passive membrane properties associated with synchronized seizure activity in L5 mPFC neurons

To directly assess how seizures alter the heterogeneity of intrinsic neuronal properties, we compared passive membrane properties of L5 mPFC pyramidal neurons in kainic acid-injected (KA, seizure-prone) versus saline-injected (SA, non-seizure-prone) mice using a series of hyperpolarizing current steps (–50 to –200 pA in 50 pA increments; 600 ms each; **Fig. 3A**).

**Figure 3.**
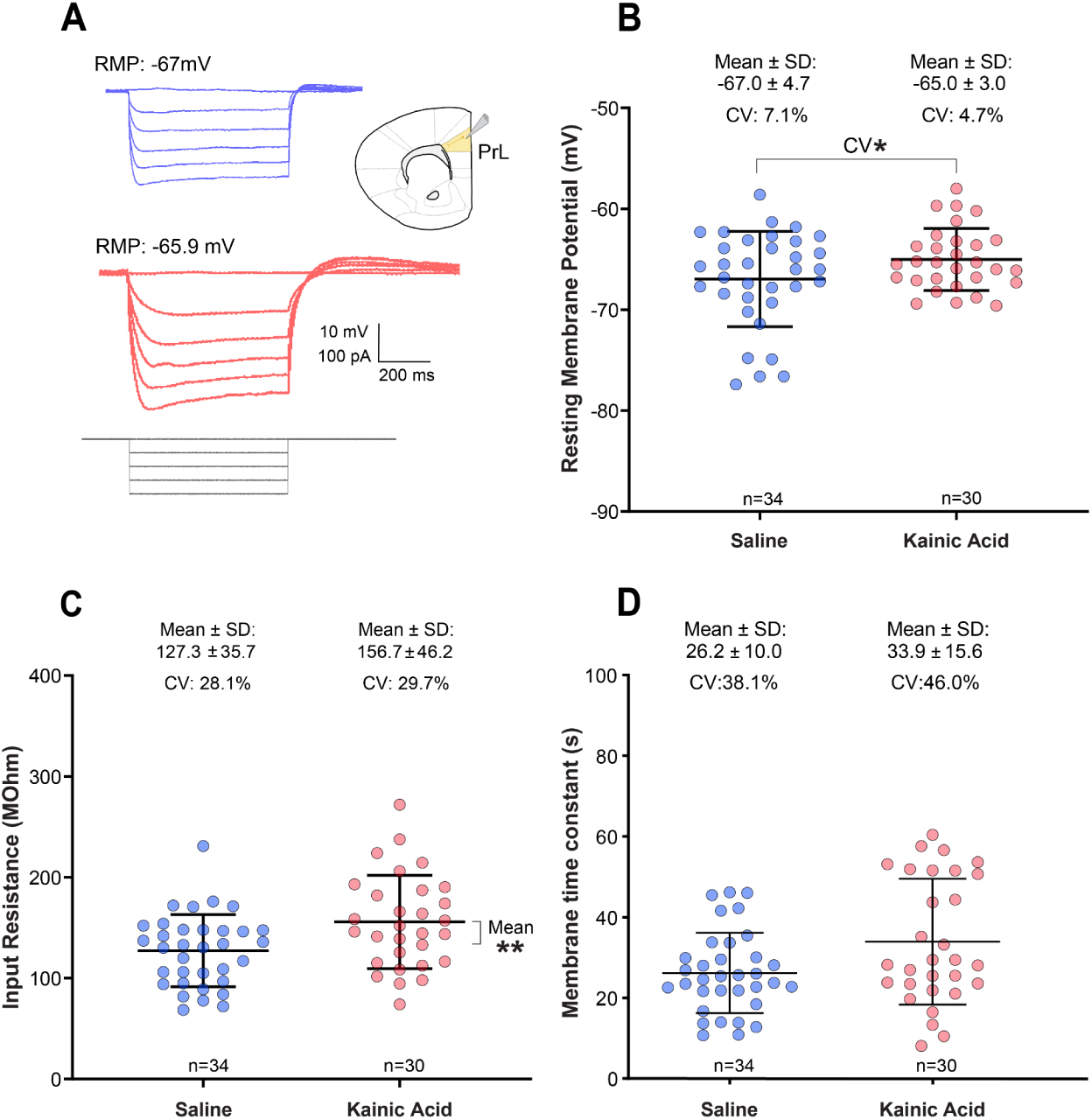
| Differences in electrophysiological properties, mean and variability, of L5 mPFC pyramidal neurons between seizure-prone (kainic acid-injected) and non-seizure-prone (saline-injected) conditions: **(A)** Schematic representation of whole-cell patch-clamp recordings targeting L5 of the mPFC and example voltage responses to hyperpolarizing current steps in saline and kainic acid conditions. **(B)** RMP in L5 mPFC neurons showed no significant difference in mean values between groups (p = 0.0542; unpaired Welch’s t-test), but variability was significantly reduced in the epileptogenic condition (p = 0.0408, two-sample coefficient of variation test). **(C)** Input resistance was significantly increased in the epileptogenic group (KA: 156 ± 46.2 MΩ, n = 30; SA: 127 ± 35.7 MΩ, n = 34; p = 0.00860; unpaired t-test with Welch’s correction), while variability remained unchanged (KA: CV = 29.7%; SA: CV = 28.1%; p = 0.281; two-sample coefficient of variation test). **(D)** Membrane time constant also showed no significant differences in either mean values (KA: 33.9 ± 15.6 ms; SA: 26.2 ± 9.97 ms; p = 0.0584; Mann-Whitney test) or variability (KA: CV = 46.0%; SA: CV = 38.1%; p = 0.369; two-sample coefficient of variation test).

RMP variability was significantly reduced in the seizure-prone group (KA: CV = 4.7%, n = 30) relative to non-seizure-prone (SA: CV = 7.1%, n = 34; p = 0.0408; **Fig. 3B**), indicating a seizure-associated decline in resting potential heterogeneity. This reduction in variability occurred without a significant change in mean RMP (KA: –65.0 ± 3.08 mV; SA: –67.0 ± 4.73 mV; p = 0.0542; unpaired t-test with Welch’s correction). In contrast, input resistance variability remained unchanged between groups (KA: CV = 29.7%; SA: CV = 28.1%; p = 0.28; **Fig. 3C**), although the mean input resistance was significantly higher in the KA group (KA: 156 ± 46.2 MΩ; SA: 127 ± 35.7 MΩ; p = 0.00860; unpaired t-test with Welch’s correction; **Fig. 3C**). Similarly, no significant differences in the variability of the membrane time constant were observed (KA: CV = 46.0%; SA: CV = 38.1%; p = 0.369; **Fig. 3D**), without a change in the mean of the time constant in Kainic acid-injected mice (KA: 33.9 ± 15.6 ms; SA: 26.2 ± 9.97 ms; p = 0.0584; Mann-Whitney test; **Fig. 3D**).

Similar to human slice cultured neurons, mPFC neurons from seizure-prone mice demonstrated a reduction in RMP heterogeneity. Notably, while overall variability in passive membrane properties remained largely intact, the isolated decrease in RMP variability may have outsized functional consequences. Prior work has shown that reduced RMP heterogeneity can destabilize network dynamics (Hass et al., 2022), suggesting that even this single change could increase the susceptibility of L5 mPFC neurons to entrain to correlated inputs arriving from the epileptogenic zone. This enhanced synchrony may facilitate the recruitment of mPFC into seizure activity and support its propagation to downstream brain regions, despite minimal changes in other passive properties.

#### 3.2.2 Reduced variability in active electrophysiological properties under epileptogenic conditions in L5 mPFC neurons

Seizure-prone conditions consistently led to reductions in intrinsic biophysical heterogeneity, even when mean values remained unchanged. In L5 mPFC pyramidal neurons, time to spike peak variability was significantly reduced at 50 and 100 pA current steps (50 pA: p = 0.0351; 100 pA: p = 0.0268; two-sample coefficient of variation test; **Fig. 4A**), despite no significant difference in the mean time to spike peak. Similarly, interspike interval variability (LV-ISI) was significantly lower in the seizure-prone group at 150 pA (KA: CV = 127.2%; SA: CV = 266.7%; p = 0.047; **Fig. 4B**), even though mean LV-ISI was unaffected. These findings point to a more temporally homogeneous firing pattern of neurons in the seizure-prone state, which can lead to easier entrainment into synchronous activity.

**Figure 4.**
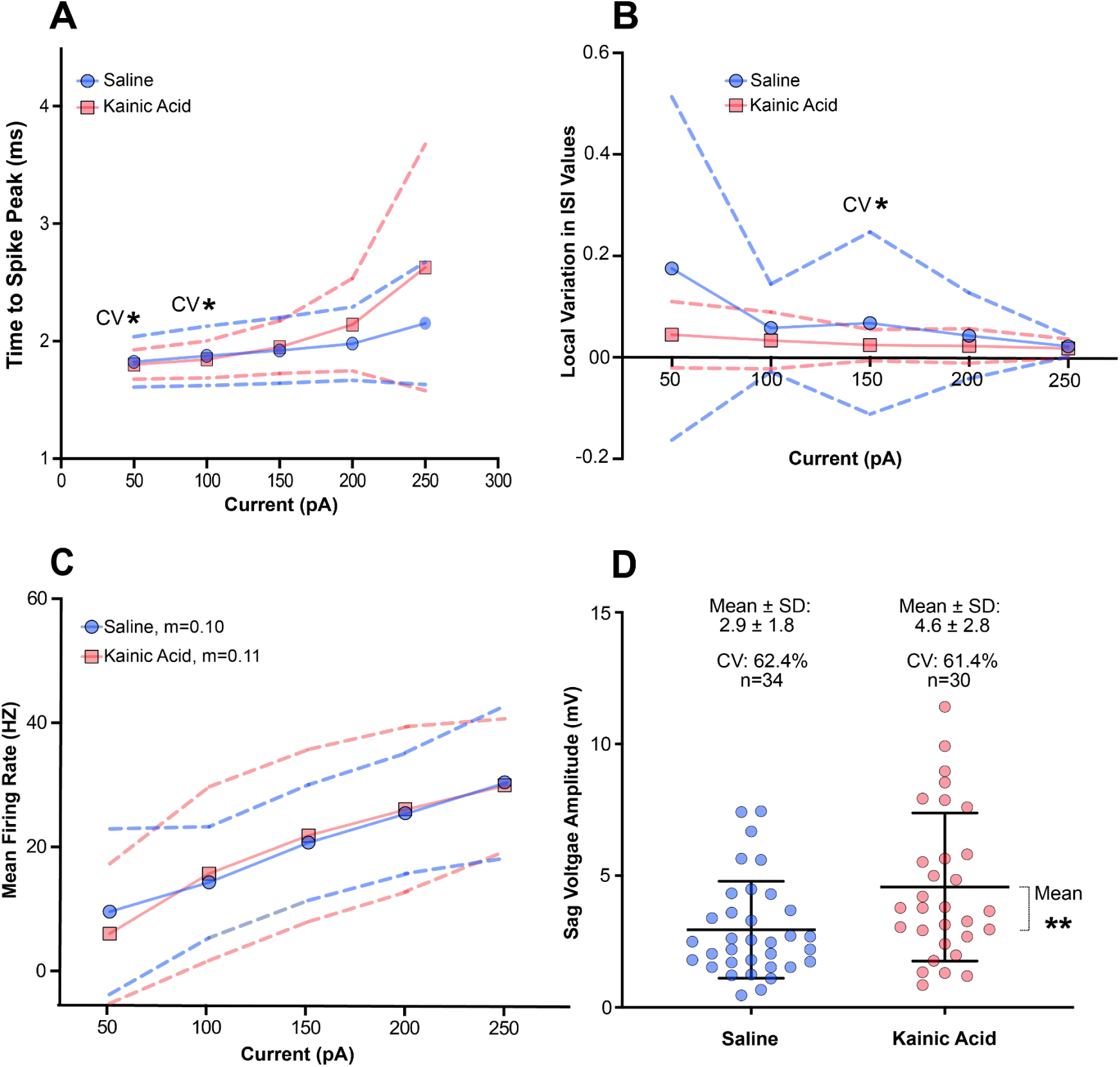
| Reduced active electrophysiological heterogeneity in L5 mPFC pyramidal neurons following seizure activity: (A) Time to spike peak was assessed across current steps, revealing unchanged mean values but significantly reduced variability in the seizure-prone group at 50 and 100 pA (CV comparison: 50 pA, p = 0.0351; 100 pA, p = 0.0268; two-sample coefficient of variation test). **(B)** Spike timing regularity was evaluated using the LV-ISI in response to 600 ms depolarizing current injections (50–250 pA). While mean LV-ISI values did not differ between groups, variability at 150 pA was significantly reduced in seizure-prone neurons (KA: CV = 127.2%) compared to non-seizure-prone (SA: CV = 266.7%; p = 0.047; two-sample coefficient of variation test), indicating increased spike timing uniformity under seizure-prone conditions. **(C)** In L5 pyramidal neurons of the mPFC, both the mean firing rate and spiking variability remained similar between seizure-prone and non-seizure-prone conditions across all current injections. No difference in the slope (m) of the firing rate was observed. **(D)** Sag voltage amplitude, a measure of HCN channel activity, was significantly higher in seizure-prone L5 mPFC pyramidal neurons (KA: 4.6 ± 2.8 mV, n = 30) than in non-seizure-prone neurons (SA: 2.9 ± 1.8 mV, n = 34; p = 0.005; Mann-Whitney U test). However, variability in sag amplitude remained comparable between groups (KA: CV = 61.4%, SA: CV = 62.4%; p = 0.947).

As previously reported (Rich et al., 2022), no significant differences were observed in either the mean or variability of firing frequency across groups (**Fig. 4C**). While variability in sag voltage amplitude, a property mediated by HCN channels, was not significantly altered (KA: CV = 60.4%; SA: CV = 62.4%; p = 0.947; **Fig. 4D**), its mean amplitude was significantly increased in neurons from seizure-prone animals compared to non-seizure-prone (KA: 4.6 ± 2.8 mV, n = 30; SA: 2.9 ± 1.8 mV, n = 34; p = 0.005; Mann-Whitney test).

To determine whether this reduction in intrinsic variability extends beyond the mPFC, we examined deep-layer pyramidal neurons in the subiculum (**Supplementary** Fig. 3), a key node to where hippocampal seizures spread to (Fei et al., 2022). In this region as well, mean values of both passive and active properties were preserved, but variability was significantly reduced in key excitability measures. Action potential threshold variability was markedly decreased (KA: CV = 5.64%; SA: CV = 12.6%; p = 0.0000446), as was LV-ISI variability (KA: CV = 74.1%; SA: CV = 157.9%; p = 0.04; two-sample coefficient of variation test; **Supplementary** Fig. 3). A summary of mean and variability changes in intrinsic biophysical properties of pyramidal neurons in the deep subiculum and L5 mPFC between seizure-prone (Kainic acid-injected) and non-seizure-prone (saline-injected) conditions is presented in **Supplementary** Fig. 2.

Together, these findings demonstrate a convergent reduction in biophysical heterogeneity across both the mPFC and subiculum, affecting multiple features of neuronal excitability. This decline in the heterogeneity of intrinsic properties may undermine the ability of local circuits to decorrelate activity, thereby increasing the likelihood of network synchronization and facilitating seizure propagation from the epileptogenic zone into the anterior nucleus of the thalamus (Fei et al., 2022) during secondary generalization of seizures.

### 3.3 Decline in spiking dynamic heterogeneity in rodent epilepsy and human slice- cultured neurons

While traditional measures such as rheobase, spike threshold, and membrane resistance provide critical insights into the intrinsic biophysical properties of neurons, white noise–based classification approach captures high-dimensional, time-dependent features of voltage responses (Padmanabhan & Urban, 2010; Tripathy et al., 2013). Therefore, the use of white noise–based classification approach is particularly well-suited to quantify both within-neuron encoding complexity and between-neuron distinctiveness, allowing detection of subtle population-level changes in functional heterogeneity.

We applied PCA-based dimensionality reduction to voltage responses evoked by frozen white noise current injections in L5 mPFC pyramidal neurons from both seizure-prone (Kainic acid-injected) and non-seizure-prone (saline-injected) conditions, as well as in L2&3 pyramidal neurons from human cortical slice cultures at acute, 24-hour, 48-hour, and 72-hour time points post-culturing (**Fig. 5A, B & F**). These principal components were then used as features for cell classification using a k-Nearest Neighbors (k-NN) algorithm **(Fig. 5C & G)**. Classification accuracy was defined as the proportion of test sweeps correctly assigned to their neuron of origin, and was evaluated as a function of the number of principal components included. In parallel, we computed two within-neuron complexity metrics, response entropy, and the number of unique encodings (a measure of pattern heterogeneity) **(Fig. 5D-5I)**, to assess whether changes in classification accuracy were driven by alterations in the dynamics between neurons. A greater number of unique encodings indicates higher information expressivity, while entropy captures how uniformly distributed these encodings are across trials.

**Figure 5.**
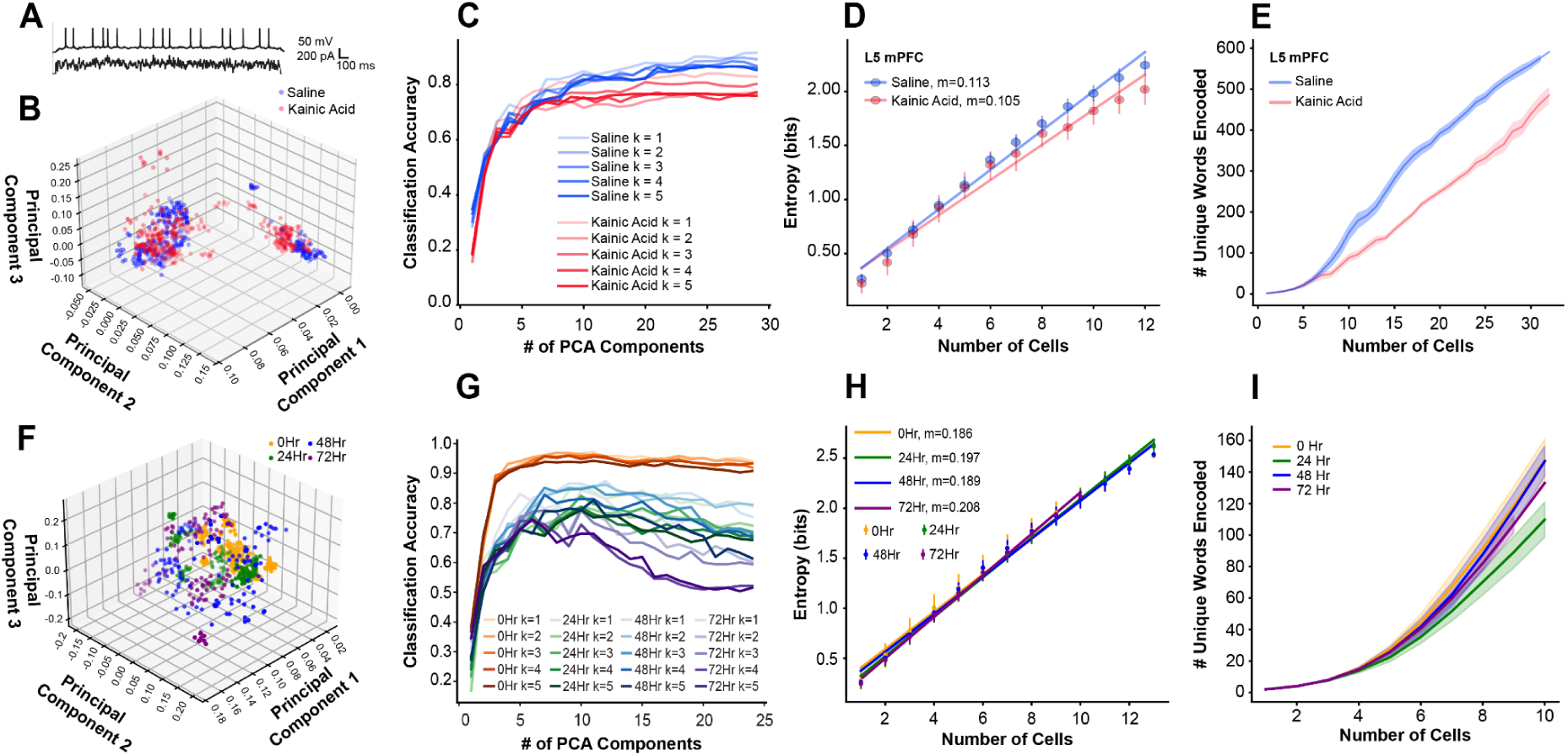
| Declines in biophysical heterogeneity are accompanied by the homogenization of spiking dynamics: **(A)** An example trace following a 2.5-second frozen white noise injection**. (B)** Following dimensionality reduction of spike trains in each treatment group with Principal Component Analysis (PCA), projection of the first 3 principal components calculated from all spike trains of all cells in control (n=33) versus seizure-prone (kainic acid-injected) (n=36) conditions**. (C)** To evaluate heterogeneity as distinctness of cells within each treatment group, we trained the k-nearest-neighbor (k-NN) algorithm on the principal components to predict the originating cell given a random spike train (averaged across different test-train splits). Our results indicate that higher prediction accuracy is achieved in the non-seizure-prone condition than in the seizure-prone condition (p<0.0001 Mann-Whitney test), meaning cells exhibit more distinctive and consistent spiking patterns. **(D-E)** There is a correlation between biophysical heterogeneity and information content metrics, such as Shannon entropy and the number of unique encodings (firing patterns) by a group of n cells, which are consistently higher in the non-seizure-prone condition compared to the seizure-prone condition for any number of cells (for entropy, control r=0.113 seizure-prone r=0.105; for number of encodings, Mann-Whitney test p<0.0001) **(D)**. Larger number of unique encodings indicates higher expressiveness, whereas higher entropy suggests a more uniform encoding distribution **(E)**. **(F)** Dimensionality reduction of spike trains in each treatment group using Principal Component Analysis (PCA), showing the projection of the first three principal components calculated from all spike trains of all cells in acute (n=13), 24-hour (n=16), 48-hour (n=13), and 72-hour (n=10) post-culturing conditions. **(G)** k-NN was performed on principal components of all 4 groups of recordings as before. The acute condition dominates post-culturing conditions in classification accuracy, followed in decreasing order of mean accuracy by 48 Hr, 24 Hr, then 72 Hr (Dunn’s test, all pairwise comparisons are significant (p<0.0001)). **(H)** Entropy remained stable across experimental groups. **(I)** In terms of the number of unique encodings, 24 Hr encodes more than 48 Hr (p<0.0001 Dunn’s test).

In the human slice culture dataset, classification accuracy was highest at early time points after slicing (acute and 24 hours), indicating strong between-neuron functional distinctiveness shortly after culturing. However, by the 72-hour time point, classification performance declined markedly which was characterized by a peak-and-fall trajectory as more PCA components were included **(Fig. 5F)**. This suggests that, at later time points, high-dimensional representations became increasingly noisy or redundant, with reduced between-neuron separability. Importantly, this reduction in classification accuracy over time occurred despite stable entropy and encoding heterogeneity (except for an increase in unique encodings at 24 hours; (p<0.0001 Dunn’s test) indicating that within-neuron encoding complexity was preserved, while between-neuron distinctions diminished **(Fig. 5H & I)**. These results suggest that in human slice cultures, intrinsic heterogeneity among neurons may erode over time in vitro, even as neuron populations maintain rich response dynamics.

In contrast, data from the K-injected mice (seizure-prone) revealed a different pattern **(Fig. 5B)**. In saline-injected animals (non-seizure-prone), classification accuracy followed a shallow incline after the initial spike (indicating the minimum number of PCs needed), steadily increasing with more PCA components and remaining high, suggesting that neuron-specific features were distributed across many dimensions and supported stable identification. However, in the seizure-prone condition, classification accuracy was significantly lower and plateaus early, indicating a loss of representation dimensions and reduction of distinguishable features **(Fig. 5B)**. Neural populations in animals displaying seizures also have reduced entropy and fewer unique encoding compared to controls **(Fig. 5E & I)**. This indicates that in the seizure-prone condition, neurons exhibited both reduced internal response complexity and less between-neuron distinctiveness, consistent with a loss of heterogeneity at both the encoding and population levels. The large number of neurons in the KA model dataset may have helped stabilize classification in control animals, allowing the classifier to benefit from even subtle between-neuron differences, while the seizure-prone condition appeared to flatten these distinctions.

Taken together, these findings demonstrate that PCA-based classification captures different underlying phenomena in the two systems: in human slice culture, a time-dependent loss of between-neuron heterogeneity without loss of individual response richness; and in the rodent KA epilepsy model, a pathology-associated compression of both within-neuron and between-neuron variability. The contrasting trajectories between the models underscore important biological differences between ex-vivo and in-vivo preparations, and between healthy and diseased brain states while highlighting that in both preparations there is heterogeneity decline in spiking dynamics.

### 3.4 HCN channel blockade restores intrinsic biophysical heterogeneity

HCN channels are heavily modulated via second messenger systems, and play a dominant role in setting a number of active and passive membrane properties of neurons. As such, our data reveals a conspicuous trend, where increases in HCN channel activity (indirectly measured from sag voltage amplitude) **(Fig. 2D and Fig. 4D)** are accompanied by decreases in excitability heterogeneity. It is likely that any ionic current that has a dominant effect on excitability, if upregulated across a neuronal population (i.e. an acquired or genetic channelopathy (van Loo & Becker, 2020)) could contribute to reduced variability in the active and passive membrane properties it regulates, although these effects are likely to be complex and nonlinear.

While our electrophysiological results suggested HCN channel upregulation, we performed genomic analysis to confirm this finding. Using whole-brain tissue collected from seizure-prone (Kainic acid-injected) and non-seizure-prone (saline-injected) mice (See “Methods”), we focused on cortical cell transcriptomics given our electrophysiological experiments. Notably, we observed upregulation of genes involved in HCN channel function, including HCN1 and its accessory subunit PEX5L (also known as TRIP8b), a key regulator of HCN channel surface expression and cyclic nucleotide sensitivity (Lewis et al., 2009, 2011) **(Fig. 6A)**. These findings suggest that upregulation of HCN channel expression contributes to the elevated sag voltage observed in L2&3 neurons of human cortical slice cultures and in L5 mPFC neurons in the KA seizure model. They also provide strong correlational evidence that regulation of HCN conductance can tune population heterogeneity.

**Figure 6.**
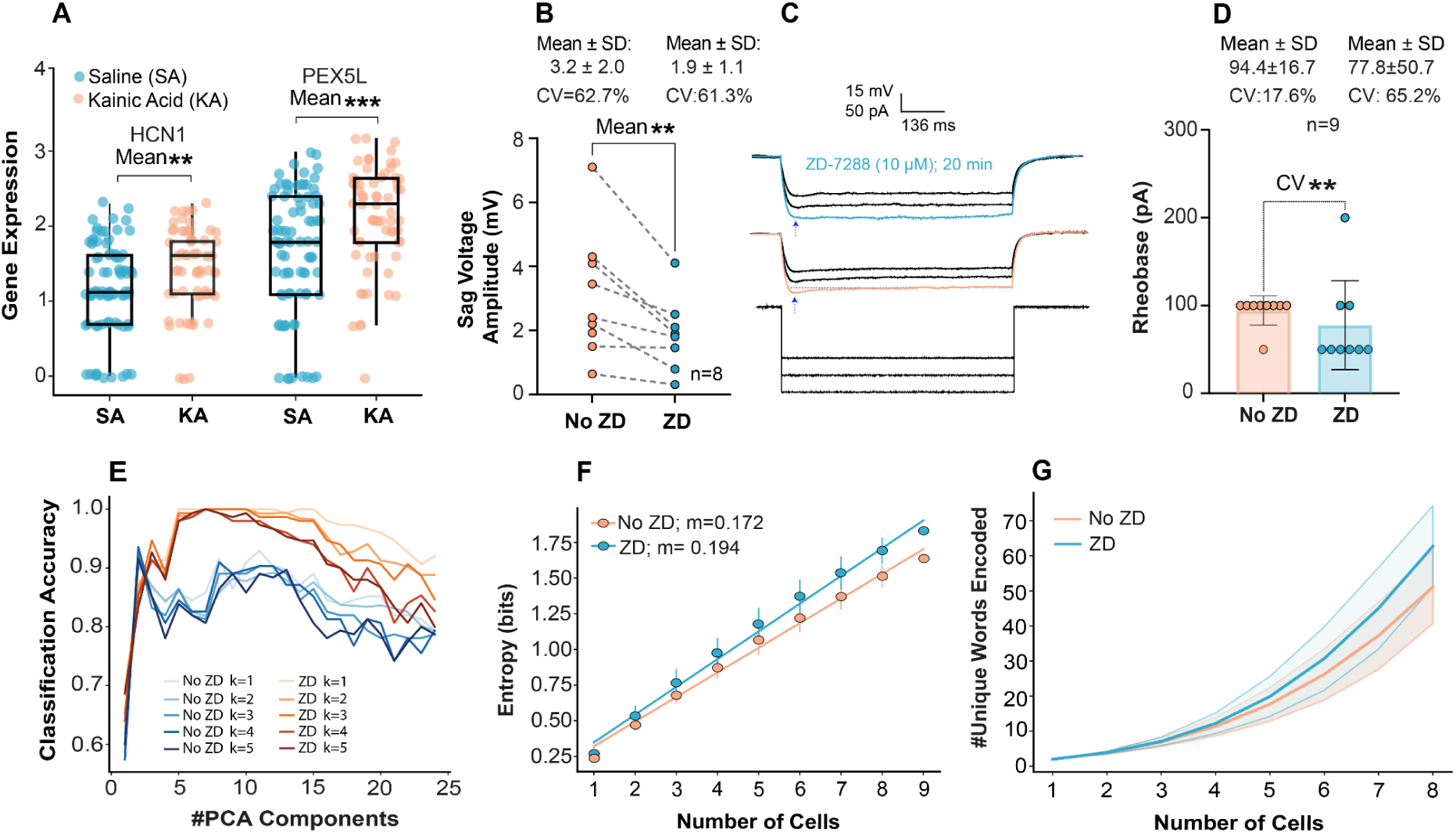
| HCN channel blockade increases electrophysiological heterogeneity: **(A)** Elevated expression of genes regulating HCN channels, including HCN1 and PEX5L (TRIP8b), was observed in whole-brain scRNA-seq data from a kainic acid-injection mouse model of temporal lobe epilepsy, compared to saline-injected controls (p < 0.05). **(B)** Whole-cell patch-clamp recordings of L5 pyramidal neurons in mouse cortex demonstrated a significant reduction in sag voltage amplitude following ZD-7288 application, confirming effective h-current blockade (**No ZD:** 3.1 ± 1.9 mV, p=0.0084, Paired t- test; CV = 63%; **ZD:** 1.9 ± 1.1 mV, CV = 61.3%; n = 8 cells, p=0.9568, two sample coefficient of variation test). **(C)** Representative voltage traces for No ZD and ZD conditions in response to a -250 pA hyperpolarizing current step are shown. **(D)** ZD-7288 application reduced the rheobase (No ZD: 94.4 ± 16.7 pA, ZD: 77.8 ± 50.7 pA, p = 0.5312; Wilcoxon matched-pairs signed rank test, n = 9) and markedly increased the heterogeneity of rheobase values across cells (No ZD, CV: 17.6%; ZD, CV: 65.2%; p = 0.0050; two sample coefficient of variation test, n = 9). **(E)** A k-NN algorithm was used to predict principal components of the recordings, revealing a significant improvement in prediction accuracy following ZD-7288 application (p < 0.0001, Mann-Whitney), indicating increased distinguishability between neurons. **(F)** Entropy analysis revealed that as the number of cells in a population increased, entropy rose linearly in both groups, but the rate of increase was faster after ZD-7288 application (No ZD: r = 0.172; ZD: r = 0.194). Mean entropy was higher after ZD-7288 application, indicating greater heterogeneity in the distribution of encodings. **(G)** The number of unique words encoded by the population of neurons was higher after ZD-7288 application (p = 0.0183, Mann-Whitney), suggesting that ZD-7288 application increased the capacity for diverse information encoding.

#### Partial restoration of electrophysiological heterogeneity following HCN channel blockade

To investigate the causal relationship between HCN channel activity and biophysical heterogeneity, we applied the HCN channel blocker ZD-7288 **(Fig. 6C)**. If increased HCN channel function reduces heterogeneity in intrinsic biophysical properties, then its blockade would be expected to restore it. Application of ZD-7288 (10 µM) significantly reduced sag voltage amplitude (*No ZD:* 3.1 ± 1.9 mV; *ZD:* 1.9 ± 1.1 mV; p = 0.0084, paired t-test), confirming effective h-current blockade, while variability (CV) remained unchanged (*No ZD:* 63%; *ZD:* 61.3%; p = 0.9568; n = 8 cells) **(Fig. 6B)**. Notably, ZD-7288 also increased the heterogeneity of rheobase across neurons (*No ZD* CV: 17.7%; *ZD* CV: 65.2%; p = 0.0050), while the change in mean rheobase was not significant (*No ZD:* 94.4 ± 16.7 pA; *ZD:* 77.8 ± 50.7 pA; p = 0.5312; Wilcoxon matched-pairs signed rank test; **Fig. 6D**). Rheobase heterogeneity has been shown to powerfully stabilize cortical dynamics (Hass et al., 2022).

Although rheobase was the only individual feature to show a statistically significant increase following ZD treatment, a broader trend toward increased variability emerged across the various other electrophysiological features (**Fig. 6D** and **Supplementary** Fig. 4). At the group level (six features in total, see **Fig. 6D** and **Supplementary** Fig. 4) we observed a statistically significant increase of 75% in the CV following ZD treatment (Wilcoxon signed-rank test on the normalized values: 1.75; *p* = 0.031). To account for the wide range and differing units of the electrophysiological features, we normalized CVs by dividing each post-treatment value by its corresponding pre-treatment value. This approach yielded unitless ratios that reflect relative changes in variability.

Moreover, ZD-7288 significantly enhanced the distinguishability of neuronal responses to frozen white noise inputs, indicating a broader spread in intrinsic electrophysiological profiles post-HCN blockade (p < 0.0001; Mann-Whitney; **Fig. 6E**). Additionally entropy analysis revealed that as the number of cells in a population increased, entropy rose linearly in both groups. However, the rate of increase was faster after the ZD-7288 application (No ZD: r = 0.172; ZD: r = 0.194). Following ZD-7288 application, mean entropy was higher, indicating greater variability in the distribution of encodings **(Fig. 6F)**. Furthermore, neurons exhibited a significantly greater capacity for information encoding after ZD-7288 application (p = 0.0183, Mann-Whitney), reflecting an increase in heterogeneity within their spike trains **(Fig. 6G)**.

These data reveal that decreasing HCN conductance levels can tune excitability variability up and increase information coding via changes in spiking dynamics. More generally these results provide the first evidence of a general mechanism by which population heterogeneity and information coding ability can be modulated by a single ionic conductance.

## 4. Discussion

Guided by our previous work demonstrating a loss of intrinsic biophysical heterogeneity in pyramidal neurons from human epileptogenic cortex (Rich et al., 2022), we explored the hypothesis that neurons with a shared history of activity will undergo a loss of biophysical heterogeneity via intrinsic plasticity mechanisms. While previous work has shown that input heterogeneity drives transcriptomic heterogeneity (Park et al., 2014), our investigation into the loss of biophysical heterogeneity is especially relevant, given its established role in predisposing neuronal networks to pathological dynamics and reducing network resilience to insults (Hutt et al., 2023; Rich et al., 2022).

We observed a marked, time-dependent reduction in within-cell heterogeneity of both passive and active intrinsic biophysical properties in cultured neurons, beginning within the first 24 hours and continuing through 72 hours post-culturing. This decline occurred over the same time scale during which intrinsic plasticity is known to operate (Debanne et al., 2019; Zhang & Linden, 2003), suggesting that a shared history of activity, imposed here via activity deprivation, is able to drive a profound loss of intrinsic biophysical heterogeneity. Concurrently, we found a progressive increase in sag voltage and a depolarization of the resting membrane potential, consistent with prior work (Ting et al., 2018). These changes are in line with previous studies showing that activity-dependent modulation of ion channel conductances serves as a homeostatic response to network quiescence (Desai et al., 1999; Johnson & Buonomano, 2007). Notably, our findings parallel those from olfactory mitral cells, where HCN channel conductance in part underlies biophysical heterogeneity (Angelo & Margrie, 2011). Crucially, we extend these insights by demonstrating that such heterogeneity is not static, rather it is a malleable property of networks that is inversely correlated with HCN channel conductance levels.

Our findings in L2&3 human cortical pyramidal neurons demonstrate that the decline in intrinsic biophysical heterogeneity is not restricted to L5 pyramidal cells (Rich et al., 2022), but instead appears to be a general feature across multiple, and possibly all cell types within the human brain. This is likely given the ubiquitous nature of intrinsic plasticity throughout all cell-types (Debanne et al., 2019; Zhang & Linden, 2003). These observations support the concept that intrinsic plasticity mediated heterogeneity decline is operative brain and layer-wide - although the extent to which heterogeneity decline would be observed in any one region of the brain will depend on the region’s and cell-type’s input statistics (Park et al., 2014; Trotter et al., 2025).

These findings are echoed in our in vivo rodent model of epilepsy, a condition characterized by excessive neural activity and synchrony. In L5 mPFC neurons from seizure-prone mice, we observed a marked reduction in the heterogeneity of both active and passive membrane properties, specifically a decreased variance in resting membrane potential (RMP). This decreased variance was as well accompanied by an increased sag voltage. RMP homogenization has been shown to destabilize network dynamics (Hass et al., 2022), analogous to our prior findings where reduced variance in distance to threshold was observed in the epileptogenic cortex (Rich et al., 2022). Complementing these physiological changes, transcriptomic analysis of seizure-prone mice revealed upregulation of HCN channel genes (**Fig. 6A**), consistent with both our electrophysiological observations and the broader framework in which intrinsic plasticity acts via modulation of ion channel conductances (Debanne et al., 2019; Zhang & Linden, 2003).

These slice culture and KA model results reveal a conserved mechanism by which heterogeneity decline can happen despite representing opposite brain states of hypo- and hyper-activity. Although precisely why HCN channel expression increases under these different conditions would be speculative, the correlation is quite clear between increased sag voltage and decreased RMP (and other) variability. Moreover, across preparations a number of other HCN dependent electrophysiological parameters (Inibhunu et al., 2023) changes were observed: (1) mean RMP; (2) threshold; and, (3) local variation of inter-spike interval. Interestingly in the subiculum we did not see a significant change in variance in RMP and sag voltage mean, although there was a decline in variability in threshold and LV-ISI. While speculative, the paucity of significant changes in electrophysiological parameter variability in subicular neurons, may be a ceiling effect of conductance level on population variance, where in non-seizure-prone condition the sag voltage is approximately four times larger in subicular neurons as compared to L5 mPFC and L2&3 human pyramidal cells. Regardless, that an acquired channelopathy (van Loo & Becker, 2020) (in this case HCN channel) can pathologically decrease biophysical heterogeneity, while its blockade can increase it (**Fig. 6 and Supplementary** Figure 4) establishes that heterogeneity can be dynamically modulated via a single control parameter/conductance.

What role do these declines in biophysical heterogeneity have on population level activity? Computational modelling has shown that biophysical heterogeneity increases information coding, while making cortical dynamics resilient to insults (Hutt et al., 2023, 2024; Rich et al., 2022). By using white noise current stimuli and information theoretic analyses (Padmanabhan & Urban, 2010), we directly explored such population level effects. We show in both preparations that information coding declines along with the declines observed in biophysical properties. Importantly, information coding could be increased by blocking HCN channels, revealing that the conductance level of a single ion channel can tune the information coding properties - up and down - of a population. These results provide a mechanistic explanation for heterogeneity “tuning” towards efficient coding (Tripathy et al., 2013). Indeed, such tuning is necessary since absolute heterogeneity is neither beneficial nor necessary for efficient neural coding (Kuncheva & Whitaker, 2003; Marsat & Maler, 2010; Mejias & Longtin, 2012; Perez-Nieves et al., 2021; Tripathy et al., 2013), and its malleability through intrinsic plasticity would allow for the needed adaptation of biophysical heterogeneity to coding requirements.

The increase in HCN channel conductance we observe, consistently across both human slice culture and our rodent model of epilepsy, has broad relevance not only to pathological states such as epilepsy (Beck & Yaari, 2008; Surges et al., 2012) and stroke (Paz et al., 2013), but also to the natural process of aging (Guet-McCreight et al., 2023). Notably, age-related working memory deficits can be rescued by HCN channel blockade (Wang et al., 2007, 2011), implicating these channels in cognitive decline. While this previous work has largely attributed the cognitive effects of increased HCN expression to changes in intrinsic excitability (Wang et al., 2007, 2011), our findings offer a recontextualization: elevated HCN conductance reduces biophysical heterogeneity, thereby impairing information coding. While our experimental data and information theoretic analyses specifically demonstrate this (**Fig. 5** & **Fig. 6F**), more generally, any mechanism that compresses variability within a neural population will lead to a loss of information content and reduced information coding capacity (Valiante, 2025).

While we have studied the pathological nature of heterogeneity decline, there are physiological examples of such declines. For example, correlated gene expression has been shown to occur in large scale networks of the brain. Both functional magnetic resonance imaging studies (Richiardi et al., 2015), and human intracranial studies (Betzel et al., 2019) have shown that regions of the brain that oscillate together share transcriptomic profiles - hence the term correlated gene expression. While the mechanism underlying such correlated gene expression is speculated to be genetically “hard coded” (Betzel et al., 2019; Richiardi et al., 2015), our work here proposes a malleable (Eberwine & Kim, 2015) and thus adaptable form of correlated gene expression through intrinsic plasticity. Together with these human studies we propose that “neurons that oscillate together transcribe together” (Park et al., 2014; Santin & Schulz, 2019; Tyssowski et al., 2018) resulting in a gradual convergence in biophysical properties (Bomkamp et al., 2019; Tripathy et al., 2017). The extent of this biophysical convergence depending on how strongly and how often they oscillate together - in other words their connectivity and input statistics (Angelo et al., 2012; Mackler et al., 1992; Mills et al., 2018; Trotter et al., 2025; Tyssowski et al., 2018) that give rise to each neuron’s *activity milieu* (Eberwine & Kim, 2015; Trotter et al., 2025). Given that oscillatory activity is a fundamental mode of brain function (Buzsáki, 2006; Fries, 2015; Vinck et al., 2023), this convergence may lower the energetic cost of re-entering specific network states during information retrieval, effectively making them easier to access (Alejandre-García et al., 2020; Gast et al., 2024).

In summary, our findings reveal a robust and conserved mechanism by which intrinsic plasticity can modulate biophysical heterogeneity across cell types, cortical layers, brain states, and species. We demonstrate that shared activity histories in both hypoactive and hyperactive conditions, activity deprivation in slice culture and synchrony in epilepsy, converge on a shared outcome: reduced biophysical variability mediated in part by increased HCN channel conductance. This loss of heterogeneity is not merely a cellular curiosity but has meaningful consequences at the network level, diminishing information coding capacity and reducing the robustness of cortical dynamics. Crucially, our results show that biophysical heterogeneity is not fixed, it is malleable, and tunable via a single conductance. Extending beyond pathology, we propose that such malleability may underlie physiological adaptations, offering a dynamic substrate for correlating and decorrelating gene expression towards efficient coding. In this view, “neurons that oscillate together transcribe together” is an emergent property of intrinsic plasticity interacting with activity histories. Such plastic tuning of heterogeneity may offer a powerful, energy-efficient strategy for dynamically regulating information flow and access in the brain

## Acknowledgements

We are deeply grateful to our neurosurgical patients and their families for their generous consent to the use of tissue samples in this research. We also thank our volunteer students, Bamdad Bazrgar, Kevin David Ireland, Tharumatha Thillainadarajah, and Darren Zhang, for helping with monitoring seizure scores. We thank the Natural Sciences and Engineering Research Council of Canada (NSERC Grant RGPIN-2015-05936 to T.A.V.) and the Canadian Institutes of Health Research (CIHR Canada Graduate Scholarship Master’s Grant to M.F.).

## Author contributions

Conceptualization: T.A.V., H.M.C.; design of the experiments: T.A.V., H.M.C., M.F.; data collection: H.M.C., M.F., M.M.; data analysis and interpretation: H.M.C., M.F., Y.Y., K.A., S.J.T., C.S., T.A.V.; drafting the article: T.A.V., H.M.C., M.F; Y.Y.; review & editing: T.A.V., H.M.C., M.F; L.Z., J.L.

---

**Supplementary Table 1.**
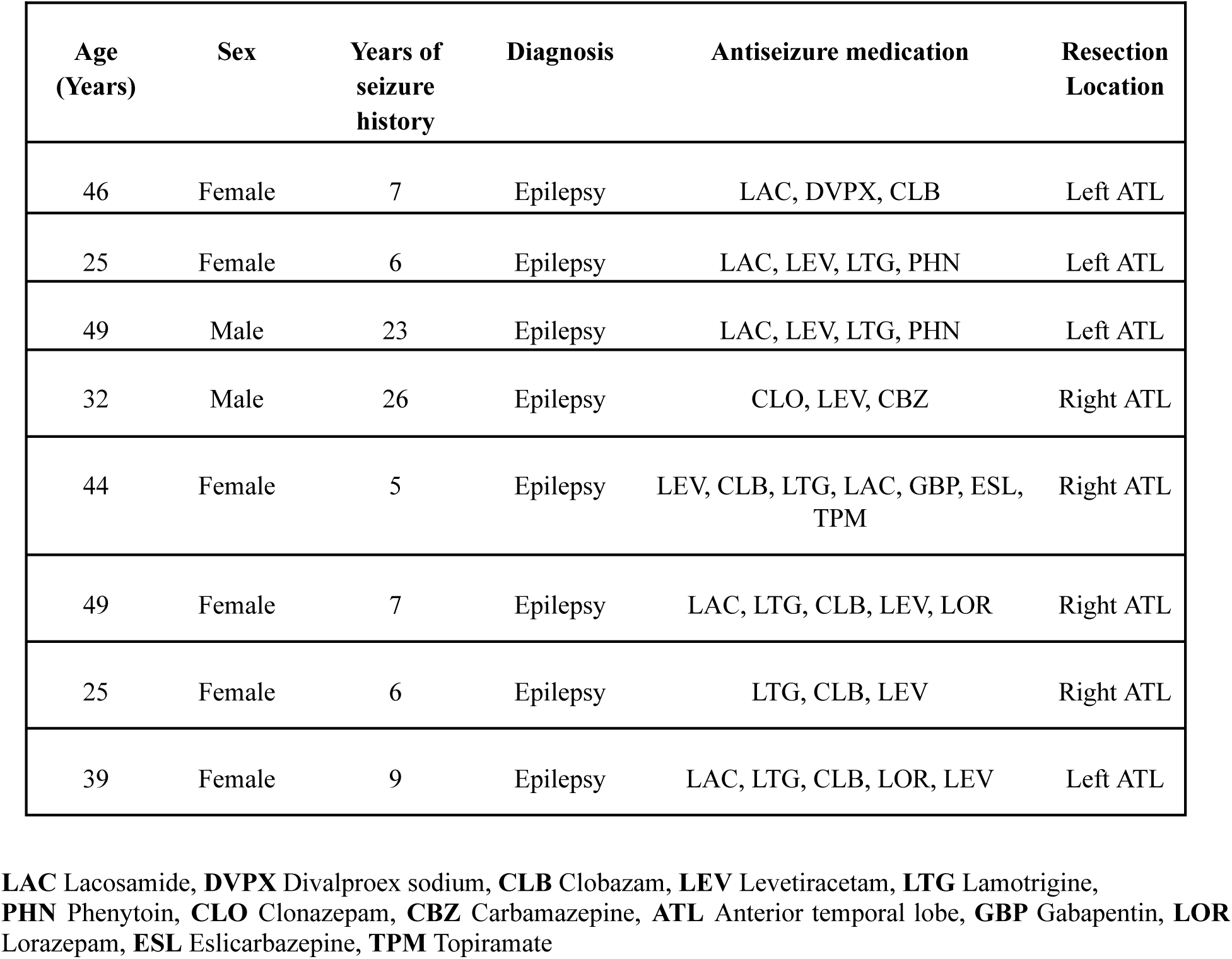
Demographic information for patients used in slice culture experiments

**Supplementary Figure 1.**
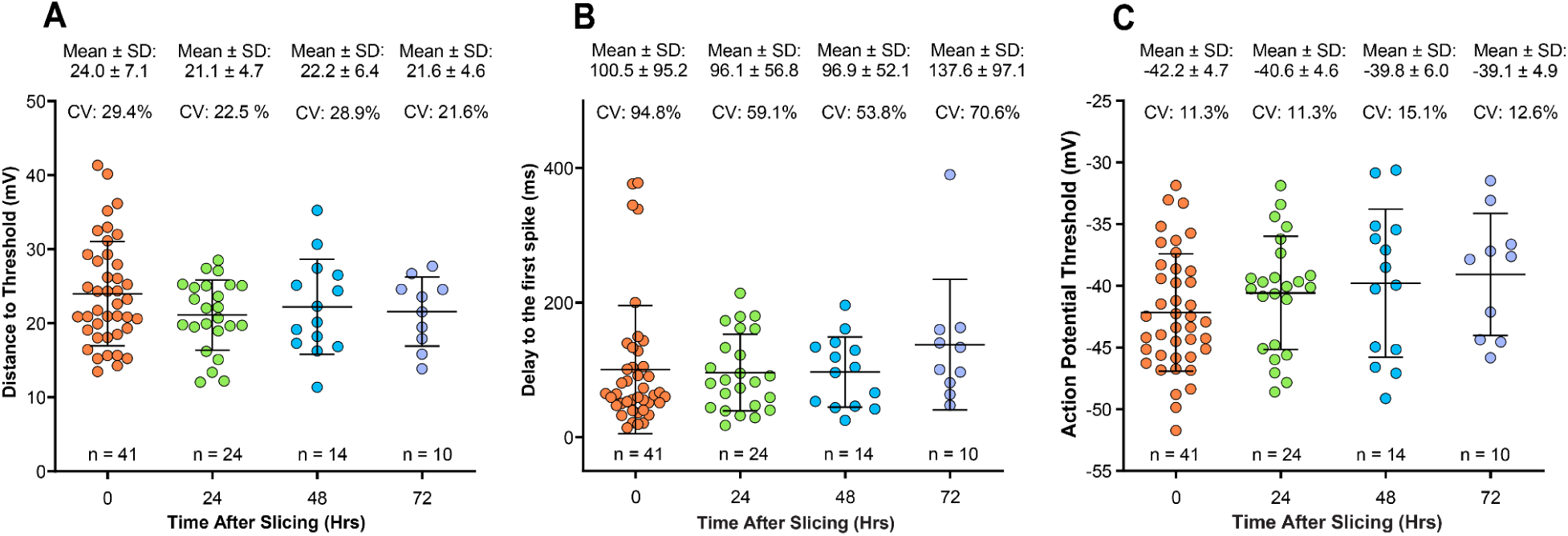
| Distance to threshold, delay to first Spike, and AP threshold remain stable in mean and variability over time in culture in L2&3 human cortical pyramidal neurons. (A) Despite the depolarization of the resting membrane potential, no significant changes were observed in the mean DTT across experimental groups (Acute: 24.0 ± 7.1 mV; 24 Hr: 21.1 ± 4.7 mV; 48 Hr: 22.2 ± 6.4 mV; 72 Hr: 21.6 ± 4.6 mV; acute Vs. 24 Hr: p = 0.2761; acute vs. 48 Hr: p = 0.7900; acute vs. 72 Hr: p = 0.6875; One-Way ANOVA followed by Tukey’s multiple comparisons test). Similarly, DTT variability remained unchanged across time points (Acute CV: 29.4%; 24 Hr CV: 22.5%; 48 Hr CV: 28.9%; 72 Hr CV: 21.6%), with no significant differences in heterogeneity compared to acute recordings (Acute vs. 24 Hr: p = 0.1927; Acute vs. 48 Hr: p = 0.9440; Acute vs. 72 Hr: p = 0.3219; Two-sample coefficients of variation tests). (B) We also quantified the delay to the first spike across experimental groups (Acute: 100.5 ± 95.2 ms; 24 Hr: 96.1 ± 56.8 ms; 48 Hr: 96.9 ± 52.1 ms; 72 Hr: 137.6 ± 97.1 ms). No significant differences were observed between any of the groups (Acute vs. 24 Hr: p > 0.9999; Acute vs. 48 Hr: p > 0.9999; Acute vs. 72 Hr: p = 0.2752; Kruskal–Wallis test followed by Dunn’s multiple comparisons test). Similarly, no significant differences were found in the variability of this measure across time points (CV: Acute = 94.8%; 24 Hr = 59.1%; 48 Hr = 53.8%; 72 Hr = 70.6%), with all comparisons to acute being non-significant (Acute vs. 24 Hr: p = 0.3571; Acute vs. 48 Hr: p = 0.2800; Acute vs. 72 Hr: p = 0.1905; two-sample coefficient of variation tests). (C)To further assess neuronal excitability, we quantified action potential threshold values across time points (Acute: –42.2 ± 4.7 mV; 24 Hr: –40.6 ± 4.6 mV; 48 Hr: –39.8 ± 6.0 mV; 72 Hr: –39.1 ± 4.9 mV). No significant differences in mean threshold were observed between any of the groups (Acute vs. 24 Hr: p = 0.5888; Acute vs. 48 Hr: p = 0.4047; Acute vs. 72 Hr: p = 0.2909; One-way ANOVA followed by Tukey’s multiple comparisons test). Similarly, there were no significant changes in variability across time points, as indicated by the coefficient of variation (Acute = 11.3%; 24 Hr = 11.3%; 48 Hr = 15.1%; 72 Hr = 12.6%; Acute vs. 24 Hr: p = 0.7836; Acute vs. 48 Hr: p = 0.1727; Acute vs. 72 Hr: p = 0.6529; two-sample coefficient of variation test).

**Supplementary Figure 2.**
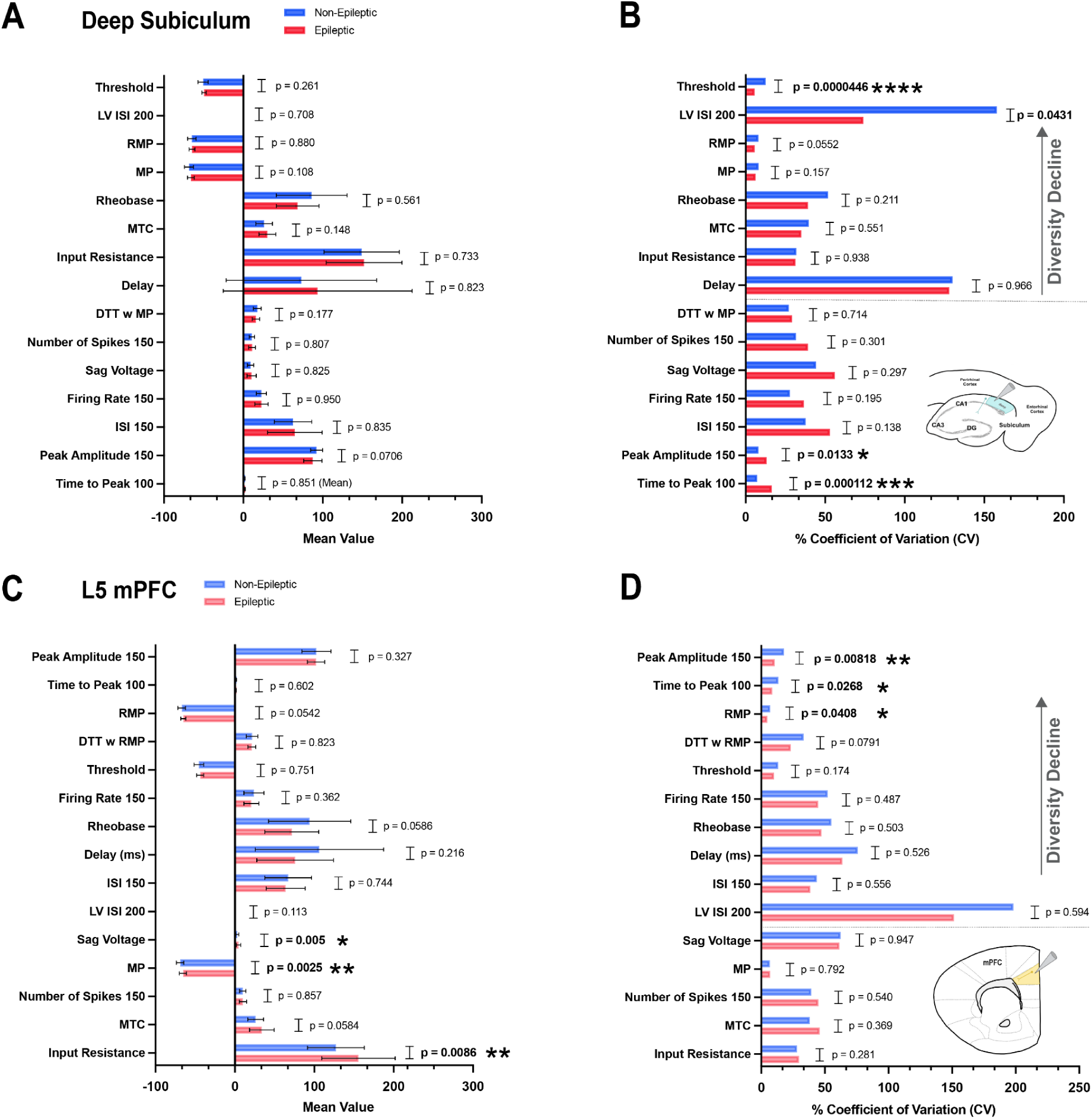
| Summary of the mean and variability changes in the intrinsic biophysical properties in the pyramidal neurons of the deep subiculum (A, B) and L5 mPFC (C, D) between the seizure-prone (kainic acid-injected) and non-seizure-prone (saline-injected) conditions. (A, C) Intrinsic biophysical properties were assessed for mean differences between conditions using either a parametric unpaired Welch’s t-test or a non-parametric unpaired Mann–Whitney test, depending on whether the data were normally or non-normally distributed. p-values for mean comparisons are displayed next to each property. (A) Deep subiculum pyramidal neurons did not show significant mean differences between the non-seizure-prone (blue) and seizure-prone (red) conditions. (B) For CV comparisons in the deep subiculum, the threshold and local variation in ISI at 200 pA showed a significant reduction in variability under seizure-prone conditions (CV differences: p = 0.0000446 and p = 0.0431, respectively). In contrast, peak amplitude at 150 pA and time to peak at 100 pA showed a significant increase in variability (CV differences: p = 0.0133 and p = 0.000112, respectively). (C) Pyramidal neurons from L5 mPFC exhibited significant mean differences in sag voltage, membrane potential (MP), and input resistance between the non-seizure-prone (light blue) and seizure-prone (light red) groups, with all measures increased in the latter. (D) Peak amplitude at 150 pA, time to peak at 100 pA, and RMP showed a significant reduction in variability, as measured by CV values, in the seizure-prone condition (CV differences: p = 0.00818, p = 0.0268, and p = 0.0408, respectively).

**Supplementary Figure 3.**
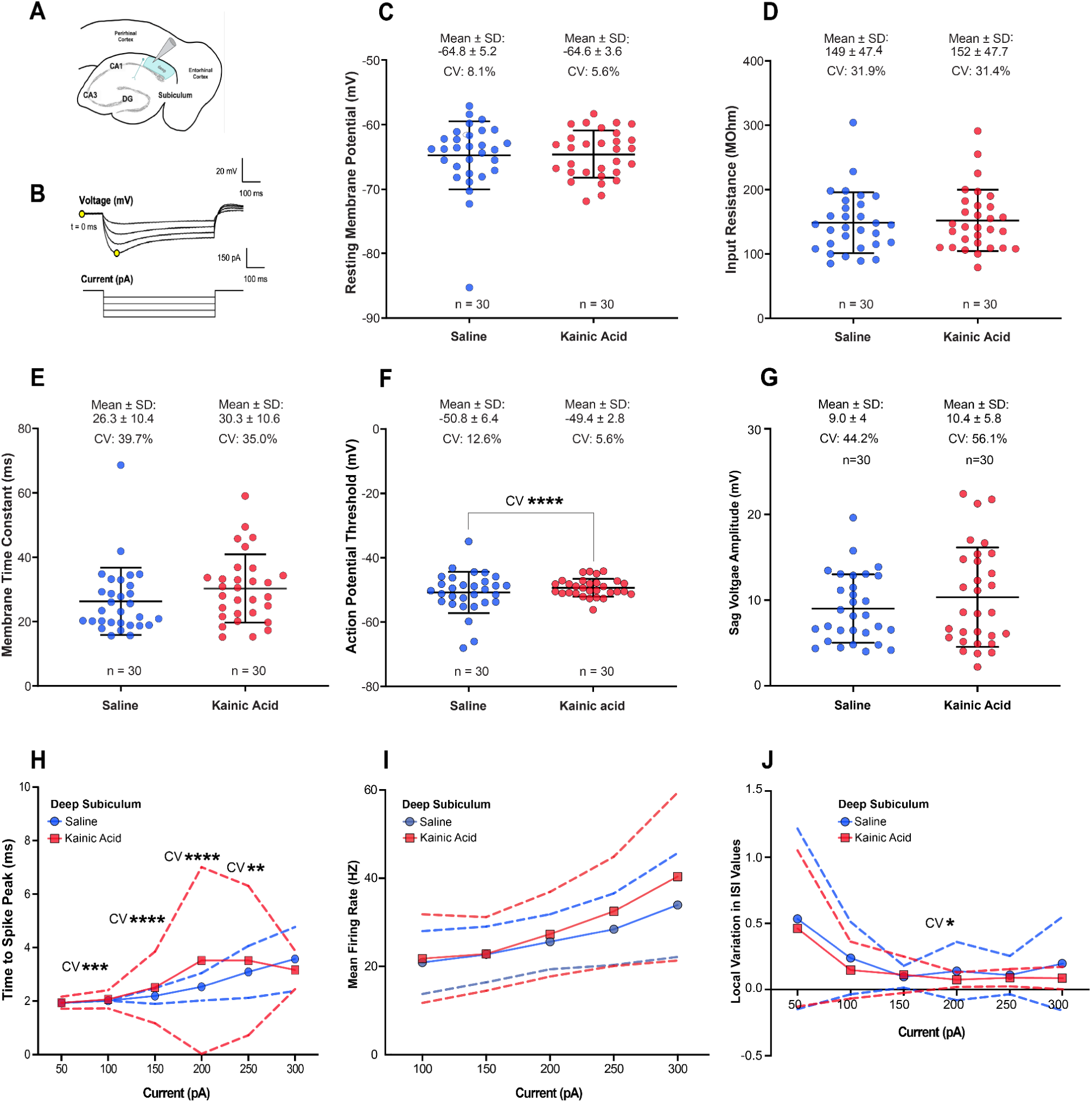
| Differences in mean and variability of electrophysiological properties in deep subiculum pyramidal neurons between seizure-prone (Kainic Acid-injected) and non-seizure-prone (Saline-injected) conditions. (A) Schematic representations of the deep subiculum regions. (B) An example of hyperpolarizing current injections and corresponding command signals used to record and calculate passive membrane properties, including membrane time constant and input resistance. (C) Resting membrane potential in deep subiculum pyramidal neurons showed no significant differences in mean (KA: -64.6 ± 3.65 mV, n = 30; saline: -64.8 ± 5.25 mV, n = 30; p = 0.880, parametric unpaired Welch’s t-test) or variability (p = 0.055, two sample coefficient of variation test, KA: CV = 5.68%; saline: CV = 8.11%) between epileptogenic and non-epileptogenic conditions. (D) Input resistance in deep subiculum neurons showed no significant differences in mean (KA: 152 ± 47.7 mOhm, n = 30; SA: 149 ± 47.4 mOhm, n = 30; p = 0.773, parametric unpaired Welch’s t-test) or variability (p = 0.525, two sample coefficient of variation test, KA: CV = 31.4%; SA: CV = 31.9%). (E) Membrane time constant in deep subiculum neurons showed no significant differences in mean (KA: 30.3 ± 10.6 ms, n = 30; saline: 26.3 ± 10.4 ms, n = 30; p = 0.148, parametric unpaired Welch’s t-test) or variability (p = 0.623, two sample coefficient of variation test, KA: CV = 35.0%; saline: CV = 39.7%) between the epileptogenic and non-epileptogenic conditions. (F) The CV values for the threshold property of deep subiculum pyramidal neurons revealed a significant reduction in variability under the epileptogenic condition (KA: CV = 12.6%) compared to non-epileptogenic condition (saline: CV = 5.64%, p = 0.0000446, two sample coefficient of variation test). This observed decline in heterogeneity was not accompanied by a significant difference in the mean values of threshold between the two conditions (KA: -49.4 ± 2.78 mV, n = 30; saline: -50.8 ± 6.41 mV, n = 30; p = 0.261, parametric unpaired Welch’s t-test). (G) Sag voltage amplitude in deep subiculum pyramidal neurons was not significantly different between seizure-prone (KA: 10.4 ± 5.8 mV, n = 30) and non-seizure-prone (SA: 9.0 ± 4 mV, n = 30; p = 0.579, parametric unpaired Welch’s t-test) conditions. Variability in sag voltage amplitude, assessed by CV values, was also not significantly different (KA: CV = 56.1%, SA: CV = 44.2%, p = 0.297). (H) Pyramidal neurons of the deep subiculum showed no significant mean differences in time to peak, measured in ms, at each current injection (50 to 300 pA). However, in terms of variability, as measured by CV, the epileptogenic condition demonstrated an increased heterogeneity from 100 pA to 300 pA compared to the non-epileptogenic condition (CV difference at 100 pA: p = 0.000112; CV difference at 150 pA: p = 2.52E-08; CV difference at 200 pA: p = 5.14E-08; CV difference at 250 pA: p = 0.000236). (I)The F-I curves illustrate the relationship between mean firing rate (Hz) and depolarizing current injections ranging from 100 to 300 pA, with shaded regions indicating standard deviation (SD). In deep subiculum pyramidal neurons, the mean firing rate did not significantly differ between seizure-prone and non-seizure-prone conditions at any current injection. (J) Local variation in inter-spike interval (LV-ISI) was analyzed to assess spike regularity in deep subiculum neurons under seizure-prone and non-seizure-prone conditions. No significant differences in mean LV-ISI were observed across current injections (50–300 pA, 600 ms). However, in the deep subiculum under epileptogenic conditions, a significant reduction in the coefficient of variation of LV-ISI was observed at 200 pA (SA: CV = 157.9%, KA: CV = 74.1%; p = 0.0431, two-sample coefficient of variation test), indicating more uniform spiking.

## Assessing intrinsic biophysical properties following partial blockade of HCN channels

Other electrophysiological properties were assessed by applying hyperpolarizing and depolarizing current injections ranging from -250 to 250 pA in 50 pA increments, with a duration of 600 ms, under both conditions: without ZD-7288 and 20 minutes after ZD-7288 application. Our results showed no significant difference in the mean RMP between the two conditions (**No ZD:** -71.0 ± 5.30 mV, n = 9; **ZD:** -68.5 ± 4.5 mV, n = 9; p = 0.2267, parametric paired t- test). Similarly, there was no significant difference in the coefficient of variation (CV) of RMP values (No ZD: CV = 7.5%; ZD: CV = 6.6%; p = 0.742; two-sample coefficient of variation test; **Supplementary** Fig 4A).

We also found no significant difference in the mean threshold between the two conditions (**No ZD:** -43.5 ± 3.4 mV, n = 9; **ZD:** -44.2 ± 5.8 mV, n = 9; p = 0.7488, paired t-test). The coefficient of variation (CV) of threshold values was also not significantly different (**No ZD:** CV

= 7.7%; **ZD:** CV = 13.2%; p = 0.1421, two-sample coefficient of variation test; **Supplementary** Fig 4B). The distance to threshold (DTT) parameter also showed no significant difference in either the mean or CV between the two conditions (**No ZD:** 27.4 ± 7.1 mV, n = 9; **ZD:** 24.3 ± 8.5 mV, n = 9; p = 0.3320, paired t-test). Likewise, there was no significant difference in the CV of threshold values (No ZD: CV = 25.8%; ZD: CV = 35%; p = 0.4295, two-sample coefficient of variation test; **Supplementary** Fig 4C). The membrane time constant was not significantly different in either the mean or CV between the two conditions (**No ZD:** 24.4 ± 8.8 ms, n = 9; **ZD:** 29.5 ± 14.2 ms, n = 9; p = 0.4042, paired t-test). Similarly, there was no significant difference in the coefficient of variation (CV) of the membrane time constant (**No ZD:** CV = 36.2%; **ZD:** CV = 48.1%; p = 0.4928, two-sample coefficient of variation test; **Supplementary** Fig 4D). Input resistance was not significantly different in either the mean or CV between the two conditions (**No ZD:** 134 ± 4 MΩ, n = 9; **ZD:** 148 ± 60.7 MΩ, n = 9; p = 0.3635, paired t-test). The coefficient of variation (CV) of input resistance also showed no significant difference (**No ZD:** CV = 26.9%; **ZD:** CV = 40.9%; p = 0.2888, two-sample coefficient of variation test; **Supplementary** Fig 4E).We also examined the effect of ZD-7288 on mean spiking firing rate and local variation in inter-spike interval (ISI), as Ih current is known to enhance the regularity of spiking. Our data show no change in either mean firing rate or the heterogeneity of spiking rates following depolarizing current injections at 50 pA (**No ZD:** 1.998 ± 5.995 Hz, n = 9; **ZD:** 5.521 ± 4.854 Hz, n = 9; p = 0.1562, Wilcoxon matched-pairs signed rank test), at 100 pA (**No ZD:** 11.4 ± 4.6 Hz, **ZD:** 15.0 ± 9.6 Hz, p = 0.1245; Paired t-test), 150 pA **(No ZD:** 20.8 ± 5.4

Hz, **ZD:** 27.1 ± 15.7 Hz, p = 0.2073; Paired t-test), 200 pA (**No ZD:** 26.9 ± 7.4 Hz, **ZD:** 29.2 ± 13.1 Hz, p = 0.4115; Paired t-test), and 250 pA (**No ZD:** 31.3 ± 9.3 Hz, **ZD:** 32.2 ± 12.31 Hz, p = 0.7096; Paired t-test) **(Supplementary** Fig 4F).

Moreover, there was no significant change in the heterogeneity of mean firing rate across conditions for each depolarizing current injection: at 50 pA (**No ZD:** CV = 300%; **ZD:** CV = 87.91%, p = 02894); 100 pA (**No ZD:** CV = 40.1%, **ZD:** CV = 64.1%, p = 0.2945), 150 pA (**No ZD:** CV = 26.1%, **ZD:** CV = 58.0%, p = 0.0651), 200 pA (No ZD: CV = 27.4%, ZD: CV = 44.9%, p = 0.2226), and 250 pA (No ZD: CV = 29.5%, ZD: CV = 38.2%, p = 0.5140). A two-sample CV test was used for comparisons between conditions at each injected current (Supplementary Fig 4F).

We also assessed local variation in inter-spike interval (LV-ISI) and did not observe any significant differences in mean or heterogeneity. Our data show no change in either the mean local variation in ISI or the heterogeneity of local variation in ISI following depolarizing current injections at 100 pA (**No ZD:** 0.015 ± 0.014, **ZD:** 0.012 ± 0.016, p = 0.1094; Wilcoxon matched-pairs signed rank test), 150 pA (**No ZD:** 0.006 ± 0.005, **ZD:** 0.042 ± 0.1, p = 0.9453; Wilcoxon matched-pairs signed rank test), 200 pA (**No ZD:** 0.009 ± 0.011, **ZD:** 0.009 ± 0.011, p

= 0.84.38; Wilcoxon matched-pairs signed rank test), and 250 pA (**No ZD:** 0.011 ± 0.010, **ZD:** 0.016 ± 0.018, p = 0.4374; paired t-test). Non-parametric Mann-Whitney tests were used for all comparisons **(Supplementary** Fig 4G).

Moreover, there was no significant change in the heterogeneity of local variation in ISI (LV-ISI) across conditions for each depolarizing current injection: 100 pA (**No ZD:** CV = 96.7%, **ZD:** CV = 132.0%, p = 0.9041), 150 pA (**No ZD:** CV = 76.7%, **ZD:** CV = 235.1%, p = 0.2222), 200 pA (**No ZD:** CV = 122.8%, **ZD:** CV = 115.2%, p = 0.9422), and 250 pA (**No ZD:** CV =

93.0%, **ZD:** CV = 111.9%, p = 0.7795). A two-sample CV test was used for comparisons between conditions at each injected current **(Supplementary** Fig 4G).

**Supplementary Figure 4.**
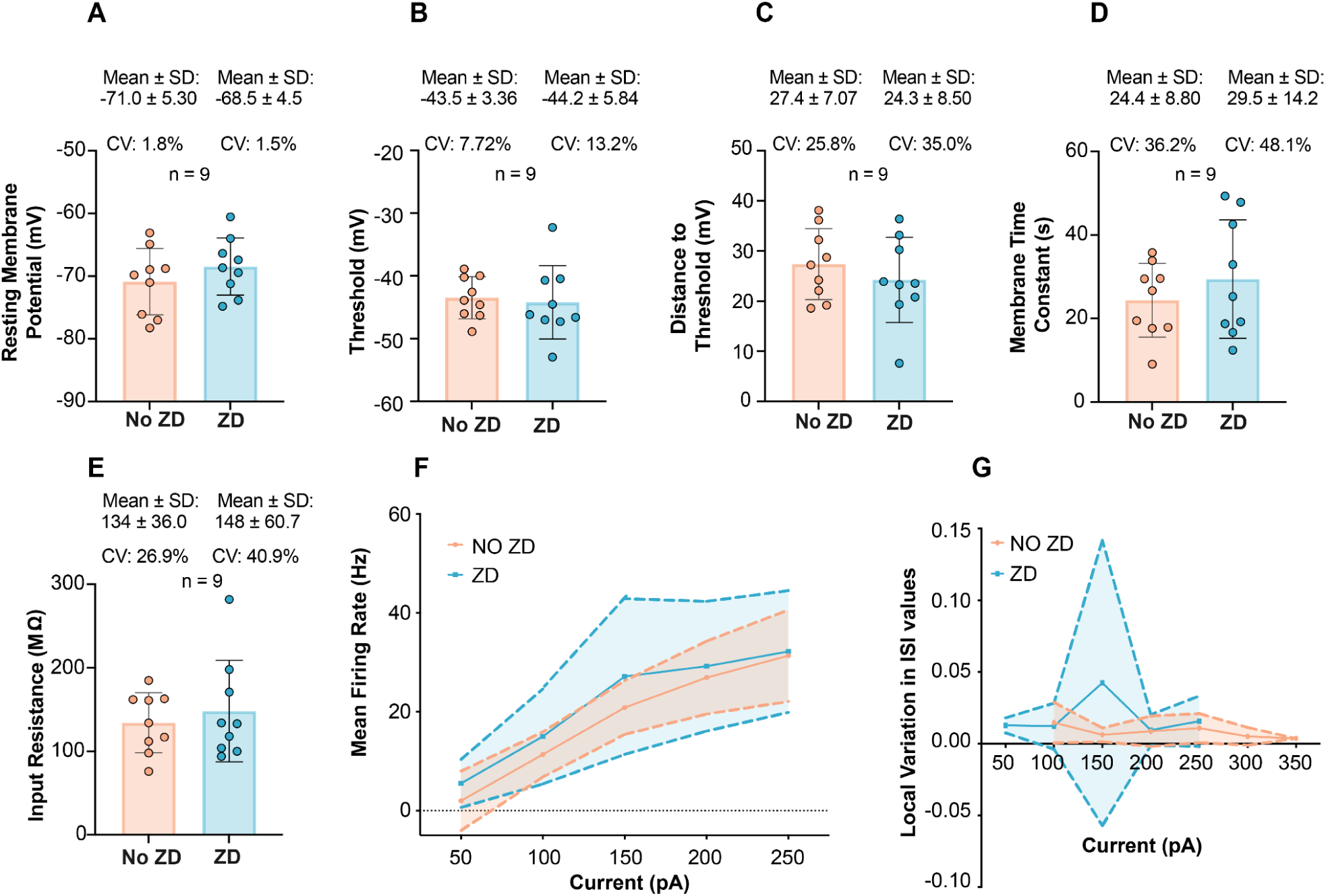
| Assessing intrinsic biophysical properties following partial blockade of HCN channels using ZD-7288 (10 µM). Passive properties such as resting membrane potential (RMP) (A), membrane time constant (MTC) (D), and input resistance (E), as well as active biophysical properties such as threshold (B), DTT (C), mean firing rate (F) and local variation in interspike interval (ISI) (G) were evaluated. Our data show no significant mean differences among these properties. However, we observed a trend toward an increase in the distribution of each parameter, though this increase was not statistically significant (No ZD, n = 9 cells; ZD, n = 9 cells).

